# Unraveling the timeline of gene expression: a pseudo-temporal trajectory analysis of single-cell RNA sequencing data

**DOI:** 10.1101/2023.05.28.542618

**Authors:** Jinming Cheng, Gordon K. Smyth, Yunshun Chen

## Abstract

There has been a rapid development in single cell RNA sequencing (scRNA-seq) technologies in recent years. Droplet-based single cell platforms such as the 10x Genomics’ Chromium system enable gene expression profiling of tens of thousands of cells per sample. The goal of a typical scRNA-seq analysis is to identify different cell subpopulations and their respective marker genes. Trajectory analysis can also be used to infer the developmental or differentiation trajectories of cells by ordering them along a putative lineage tree based on their gene expression profiles. This analysis positions cells and cell clusters along a pseudotime trajectory that represents a biological process such as cell differentiation, development, or disease progression. Here we demonstrate a time-course analysis to identify genes that are significantly associated with pseudotime. The article demonstrates a comprehensive workflow for performing trajectory inference and time course analysis on a multi-sample single cell RNA-seq experiment of the mouse mammary gland. The workflow uses open-source R software packages and covers all steps of the analysis pipeline, including quality control, doublet prediction, normalization, integration, dimension reduction, cell clustering, trajectory inference, and pseudobulk time course analysis. Sample integration and cell clustering follows the *Seurat* pipeline while the trajectory inference is conducted using the *monocle3* package. The pseudo-bulk time course analysis uses the quasi-likelihood framework of *edgeR*.

## Introduction

Single cell RNA sequencing (scRNA-seq) has emerged as a popular technique for transcriptomic profiling of samples at the single cell level. With droplet-based methods, thousands of cells can be sequenced in parallel using next-generation sequencing platforms [1, 2]. One of the most widely used droplet-based scRNA-seq technologies is the 10x Genomics Chromium which enables profiling transcriptomes of tens of thousands of cells per sample [3]. A common goal of a scRNA-seq analysis is to investigate cell types and states in heterogeneous tissues. To achieve this, various pipelines have been developed, such as *Seurat* [4] and the Bioconductor’s OSCA pipeline [5]. A typical scRNA-seq data analysis pipeline involves quality control, normalization, dimension reduction, cell clustering, and differential expression analysis.

As the cost of scRNA-seq continues to drop, more experimental studies involve replicate samples. In a multiple sample single-cell experiment, an integration method is required to investigate all cells across all samples simultaneously. This ensures that sample and batch effects are appropriately considered in visualizing and clustering cells. Popular integration methods include the Seurat’s anchor-based integration method [4], Harmony [6], and the MNN [7].

After integration and cell clustering, differential expression analysis is often performed to identify marker genes for each cell cluster. Various methods have been developed at the single-cell level for finding marker genes [8, 9]. Recently, the pseudo-bulk method has become increasingly popular due to its superior computational efficiency and its ability to consider biological variation between replicate samples [10].

Trajectory inference is another popular downstream analysis that aims to study cell differentiation or cell type development. Popular software tools to perform trajectory analysis include *monocle3* [11] and *slingshot* [12]. These methods learn trajectories based on the change of gene expression and order cells along a trajectory to obtain pseudotime [13, 14]. This allows for pseudotime-based time course analysis in single-cell experiments, which is extremely useful for investigating specific biological questions of interest.

Here we present a new single-cell workflow that integrates trajectory analysis and pseudo-bulking to execute a singlecell pseudo time course analysis. The inputs for this workflow are single-cell count matrices, such as those generated by 10x Genomic’s *cellranger* (https://www.10xgenomics.com). The single-cell level analysis is performed in *Seurat*, and the trajectory analysis is conducted using *monocle3*. Once the pseudo-bulk samples are created and assigned pseudotime, a time course analysis is conducted in *edgeR* [15]. The analysis pipeline presented in this article can be applied to any scRNA-seq study with replicate samples.

## Description of the biological experiment

The scRNA-seq data used in this workflow consists of five mouse mammary epithelium samples at five different stages: embryonic, early postnatal, pre-puberty, puberty and adult. The puberty sample is from the study in Pal et al. 2017 [16], whereas the other samples are from Pal et al. 2021 [17]. These studies examined the stage-specific single-cell profiles in order to gain insight into the early developmental stages of mammary gland epithelial lineage. The *cellranger* count matrix outputs of these five samples are available on the GEO repository as series GSE103275 and GSE164017.

## Data preparation

### Downloading the data

The *cellranger* output of each sample consists of three key files: a count matrix in `mtx.gz` format, barcode information in `tsv.gz` format and feature (or gene) information in `tsv.gz` format.

The outputs of the mouse mammary epithelium at embryonic stage (E18.5), post-natal 5 days (P5), 2.5 weeks (Prepuberty), and 10 weeks (Adult) can be downloaded from **GSE164017** [17], whereas the output of mouse mammary epithelium at 5 weeks (Puberty) can be downloaded from **GSE103275** [16].

We first create a `data` directory to store all the data files.

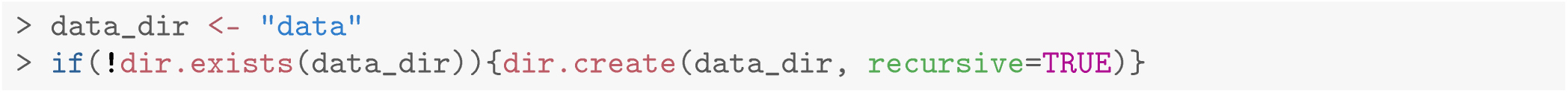

We then download the barcode and count matrix files of the five samples.

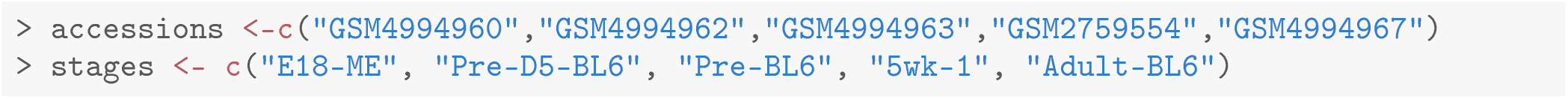

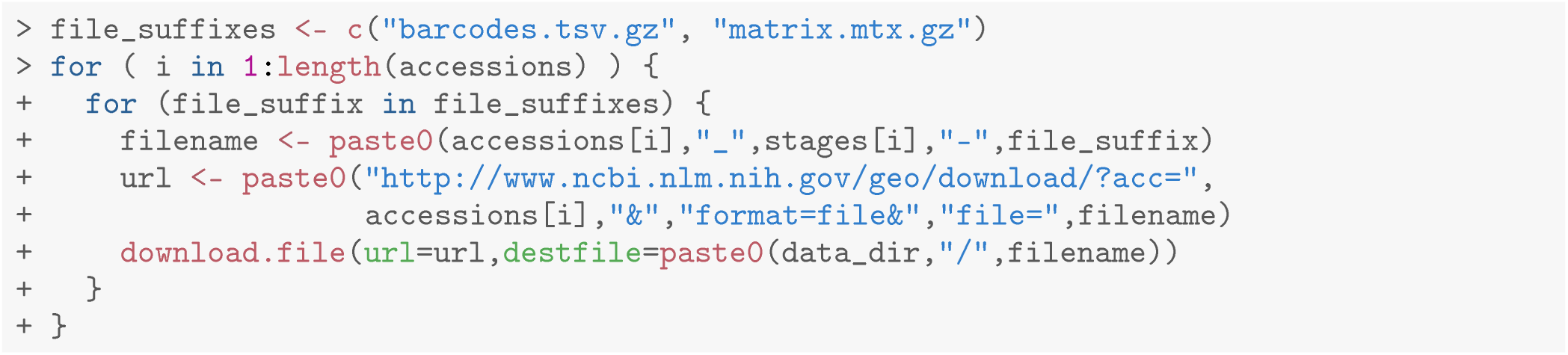

Since the five samples in this workflow are from two separate studies and were processed using different versions of mouse genome, the feature information is slightly different between the two runs. Here, we download the feature information of both runs. The `GSM2759554_5wk-1-genes.tsv.gz` file contains the feature information for the `5wk-1` sample, whereas `GSE164017_features.tsv.gz` contains the feature information for the other four samples.

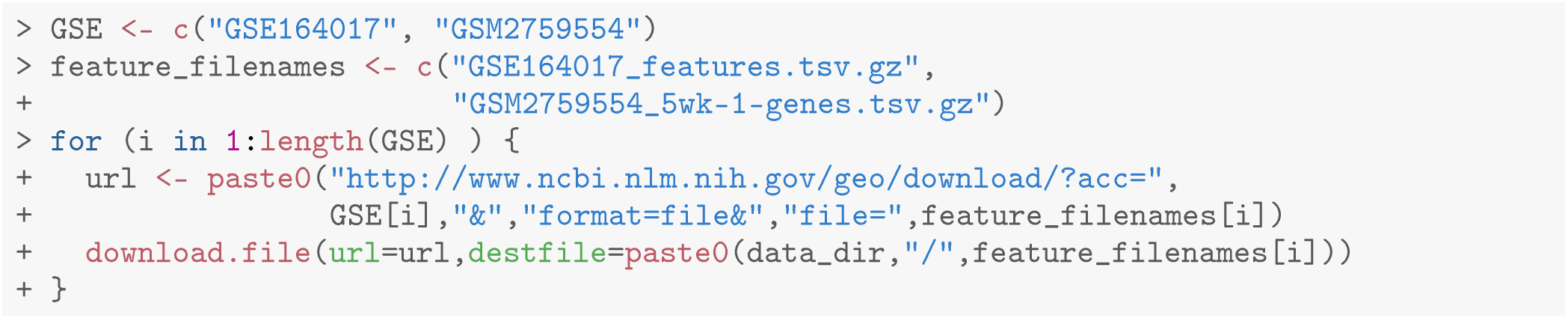

A target information file is created to store all the sample and file information.

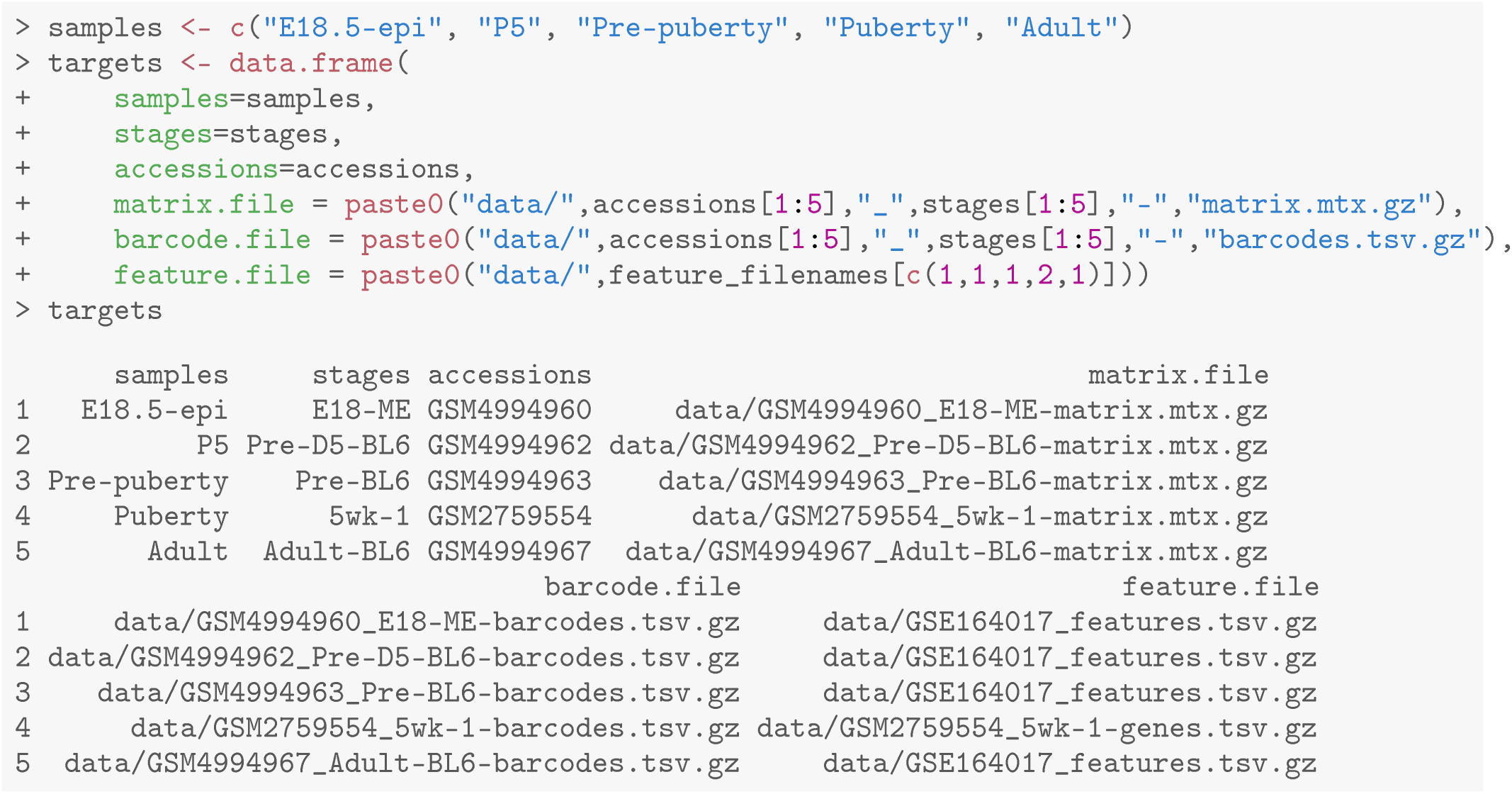

### Reading in the data

The downloaded *cellranger* outputs of all the samples can be read in one-by-one using the `read10X` function in the *edgeR* package. First, a `DGElist` object is created for each sample, which is then consolidated into a single `DGElist` object by merging them altogether.

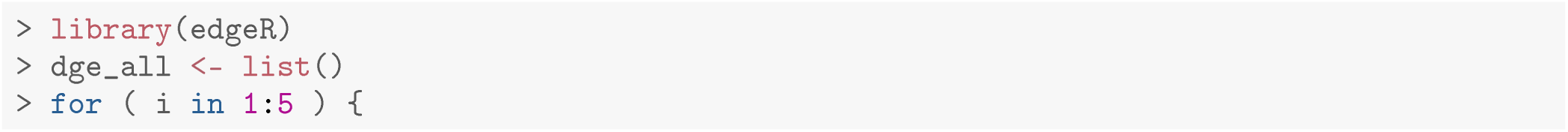

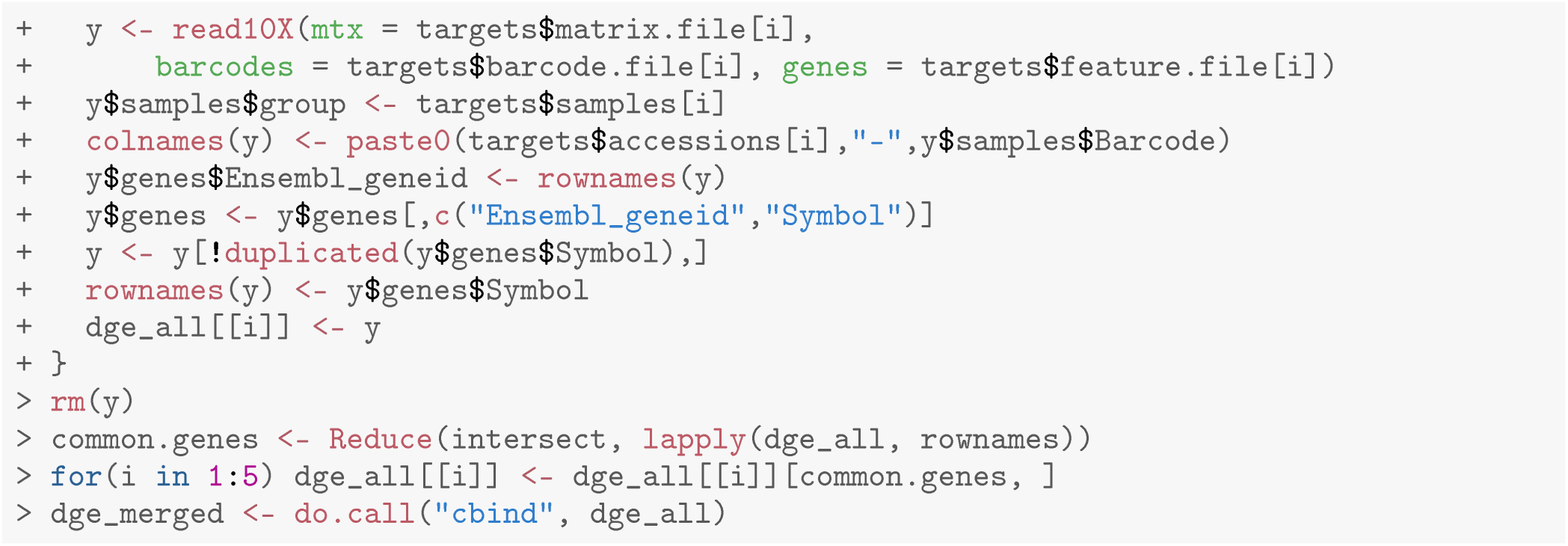

The levels of `group` in the sample information data frame are reordered and renamed from the early embryonic stage to the late adult stage.

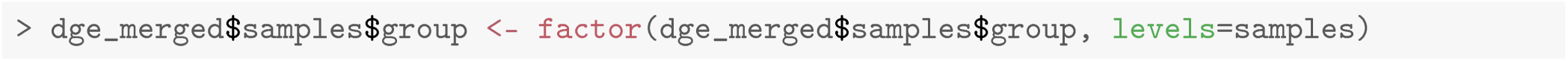

The number of genes, the total number of cells, and the number of cells in each sample are shown below.

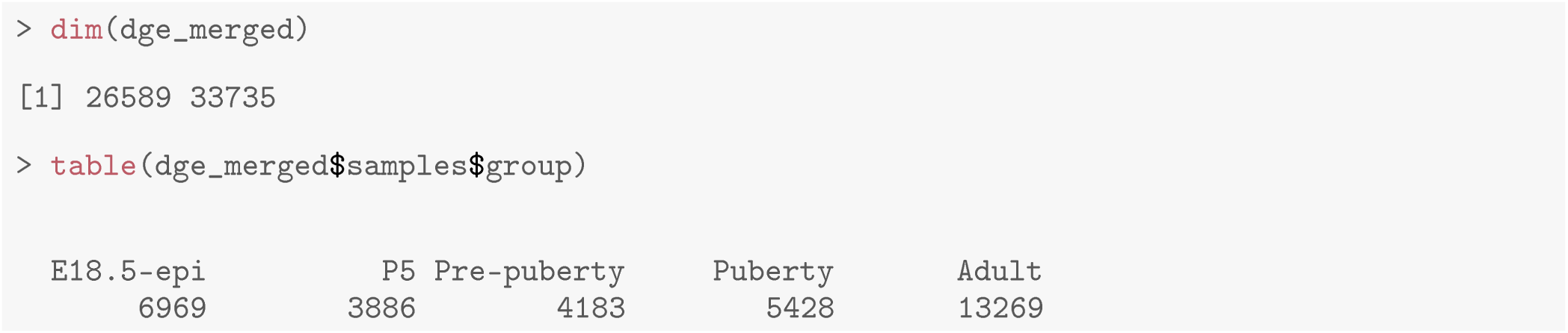

## Single-cell RNA-seq analysis

### Quality control

Quality control (QC) is essential for single cell RNA-seq data analysis. Common choices of QC metrics include number of expressed genes or features, library size, and proportion of reads mapped to mitochondrial genes in each cell. The number of expressed genes and mitochondria read percentage in each cell can be calculated as follows.

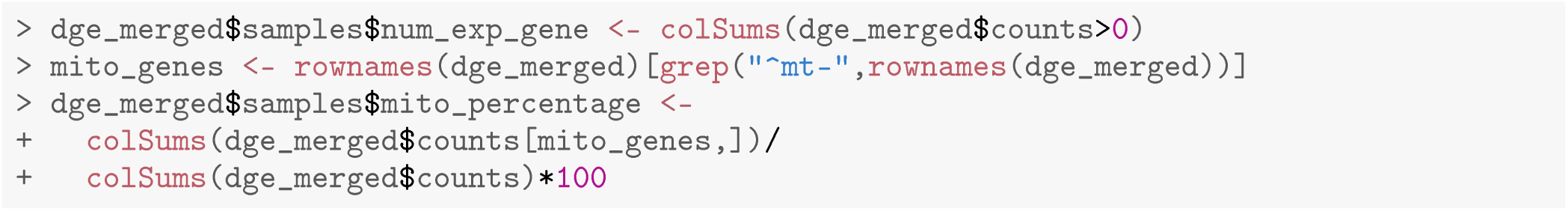

These QC metrics can be visualized in the following scatter plots (Figure 1).

**Figure 1.**
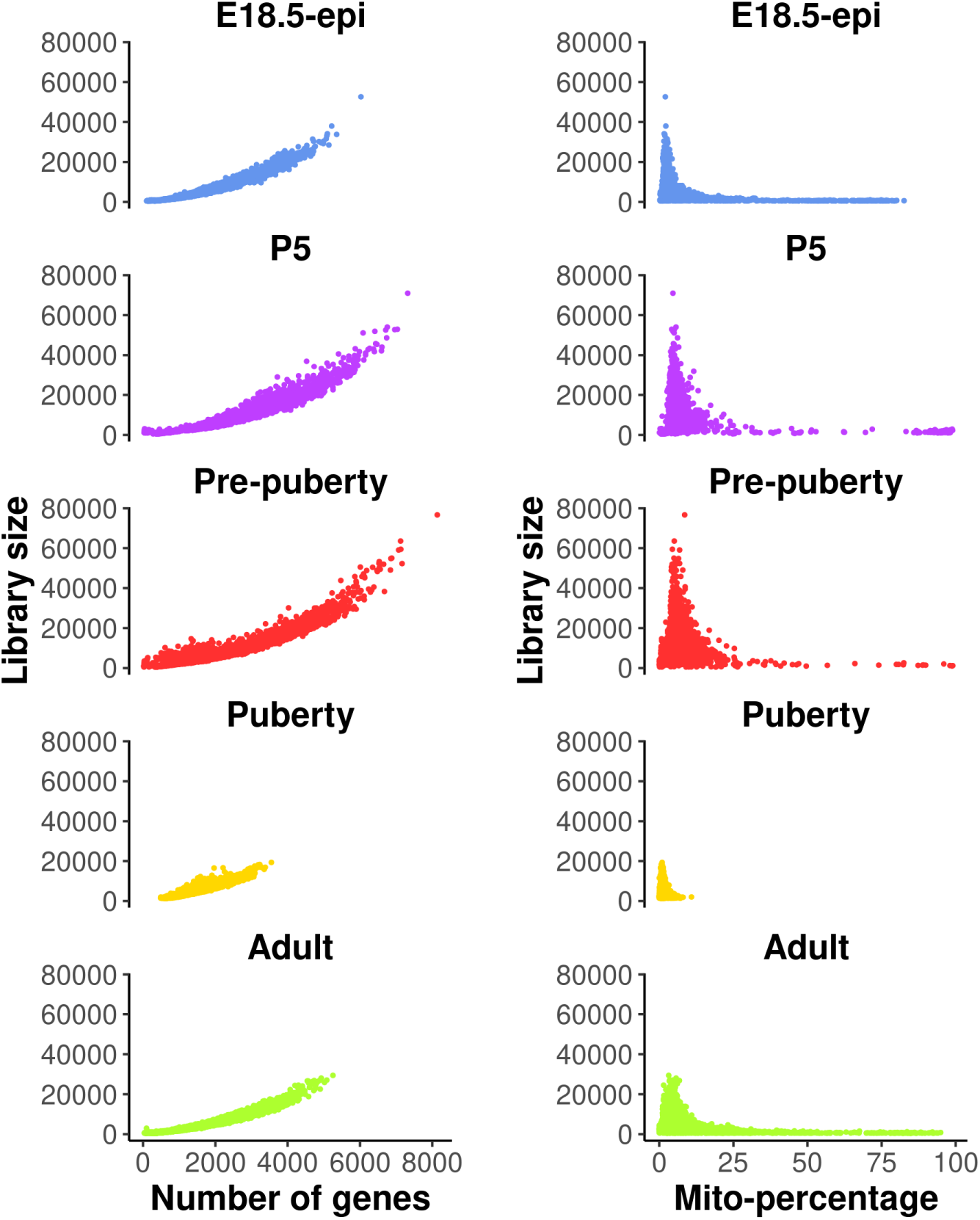
Scatter plots of quality control metrics across all the samples. The plots on the left show library size vs number of genes detected, whereas those on the right show library size vs mitochondria read percentage.

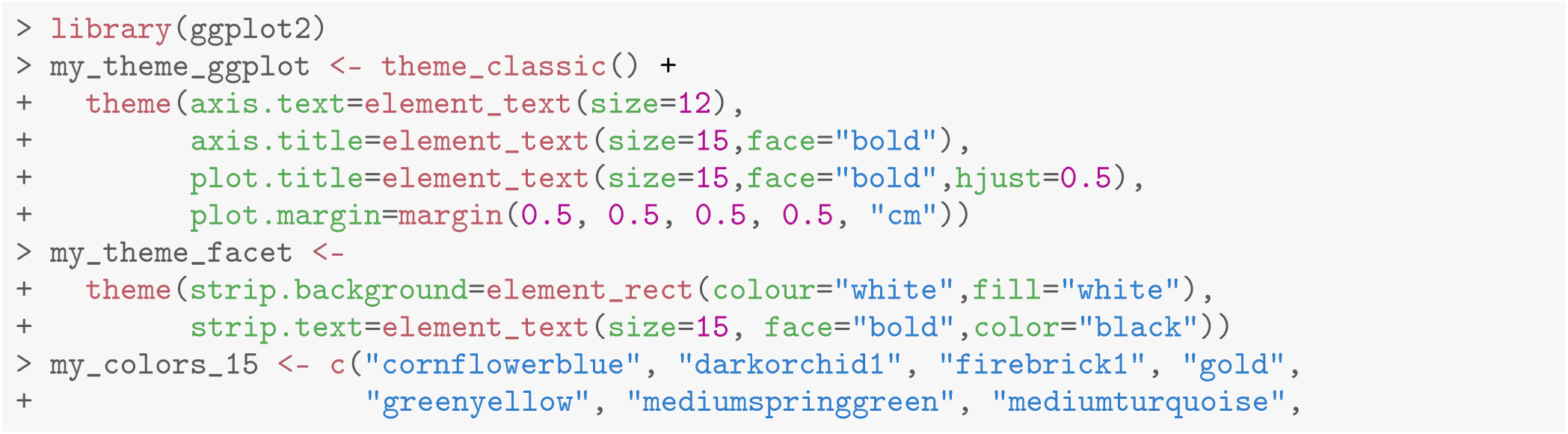

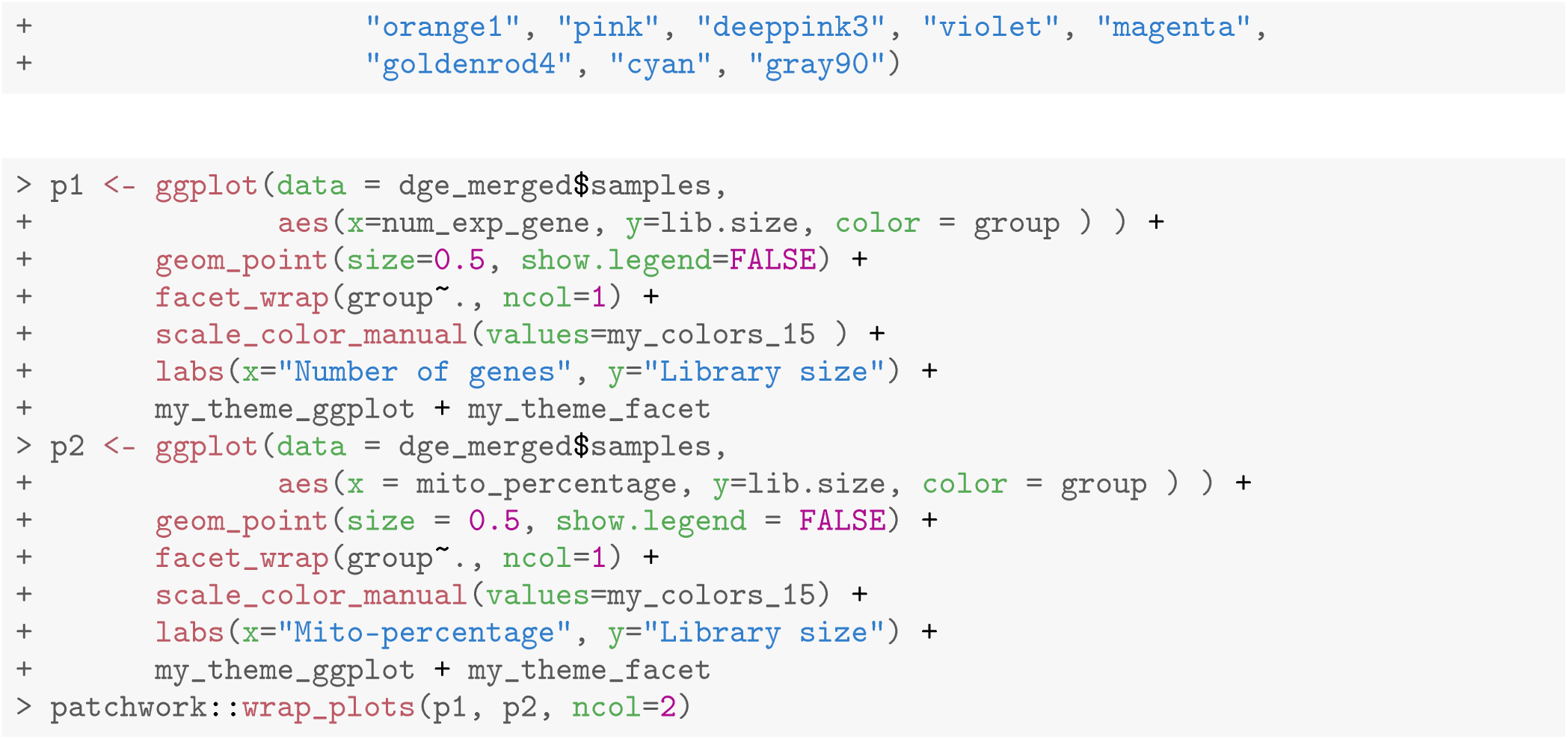

Cells with a very low number of genes (< 500), as well as high mitochondria read percentage (> 10%), are considered of low quality and hence are removed from the analysis. Cells expressing a large number of genes are also removed as they are likely to be doublets. Different thresholds are selected for different samples based on the distribution of the number of genes expressed. Here, we choose 5000, 6000, 6000, 3000, 4000 for E18.5-epi, P5, pre-puberty, puberty and adult samples, respectively. In this workflow, most of the single-cell analysis is conducted using the ***Seurat*** package. A list of five Seurat objects are first created to store the data after QC.

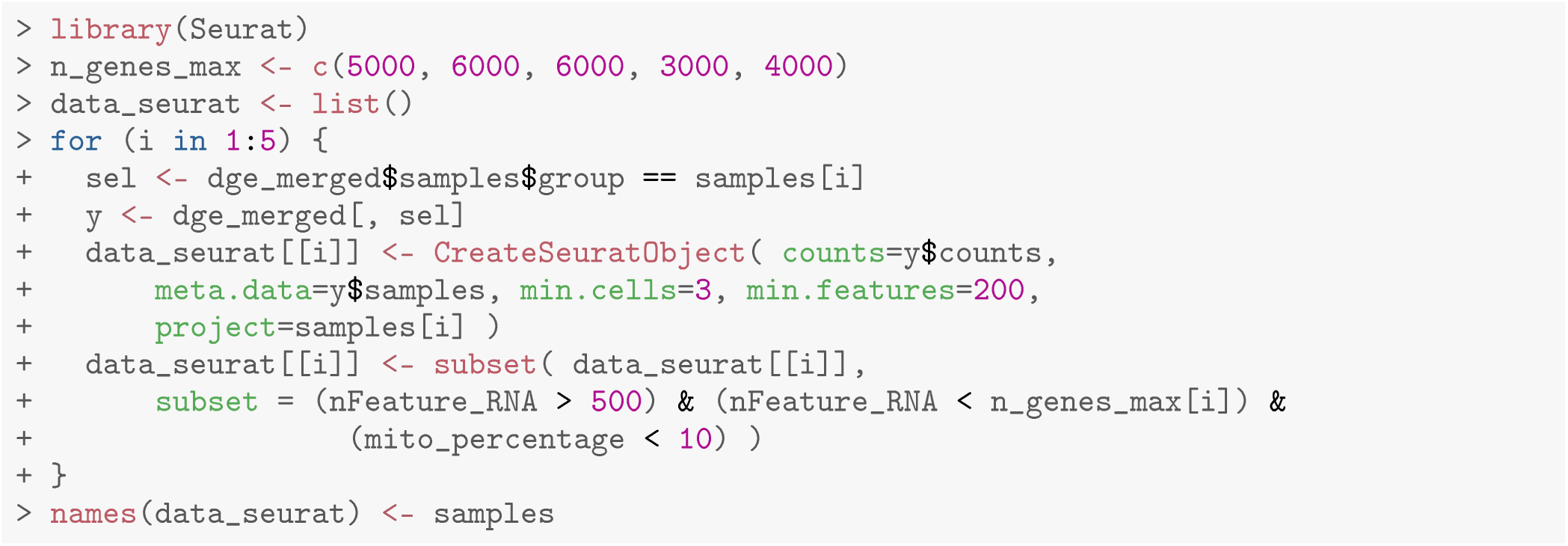

### Standard Seurat analysis of individual sample

A standard Seurat analysis is performed for each individual sample. In particular, the data of each sample is first normalized by the default log normalization method in `NormalizeData`. The top 2000 highly variable genes are identified by `FindVariableFeatures`. The normalized data of the 2000 highly variable genes are scaled by `ScaleData` to have a mean of 0 and a variance of 1. The principal component analysis (PCA) dimension reduction is performed on the highly variable genes by `RunPCA`. Uniform manifold approximation and projection (UMAP) dimension reduction is performed on the first 30 PCs by `RunUMAP`. Cell clustering is performed by `FindNeighbors` and `FindClusters`. The cell clustering resolution is set at 0.1, 0.1, 0.2, 0.2 and 0.2 for E18.5-epi, P5, pre-puberty, puberty and adult, respectively.

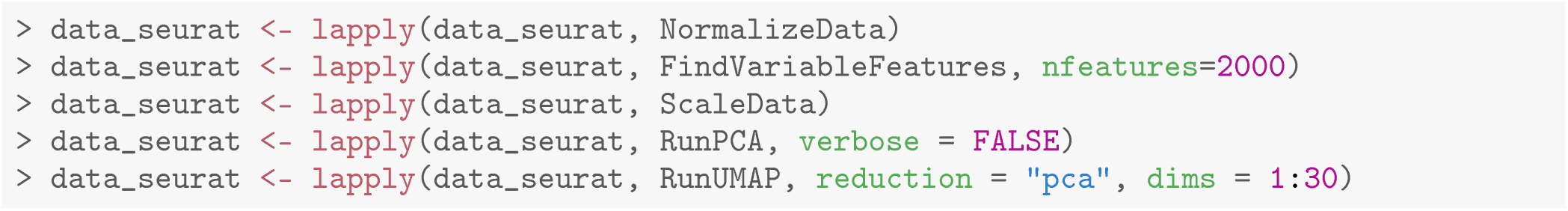

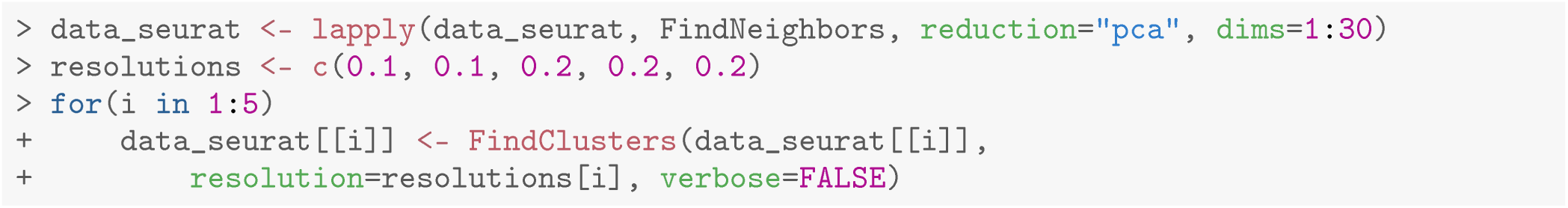

### Removing potential doublets and non-epithelial cells

Although high-throughput droplet-based single-cell technologies can accurately capture individual cells, there are instances where a single droplet may contain two or more cells, which are known as doublets or multiplets. Here we use the *scDblFinder* package [18] to further remove potential doublets. To do that, each Seurat object in the list is first converted into a `SingleCellExperiment` object using the `as.SingleCellExperiment` function in *Seurat*. Then the `scDblFinder` function in the *scDblFinder* package is called to predict potential doublets on each `SingleCellExperiment` object. The `scDblFinder` output for each sample is stored in the corresponding Seurat object.

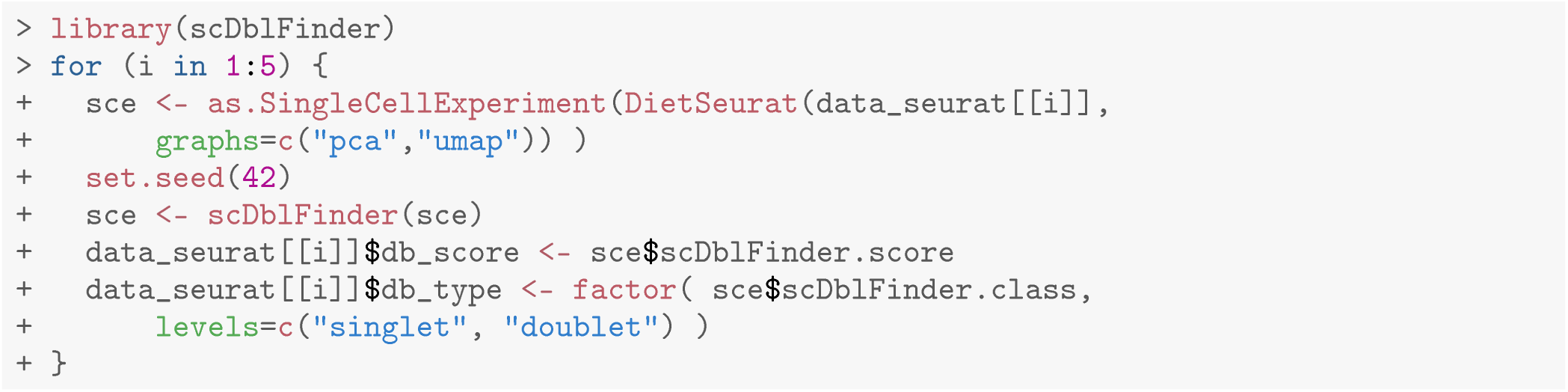

The main object of this single-cell experiment is to examine the early developmental stages of the mouse epithelial mammary gland. Therefore, we focus on epithelial cells for the rest of the analysis. We use the *Epcam* gene to identify epithelial cell clusters in each sample. The cell clustering, the expression level of *Epcam* and doublet prediction results of each sample are shown below (Figure 2).

**Figure 2.**
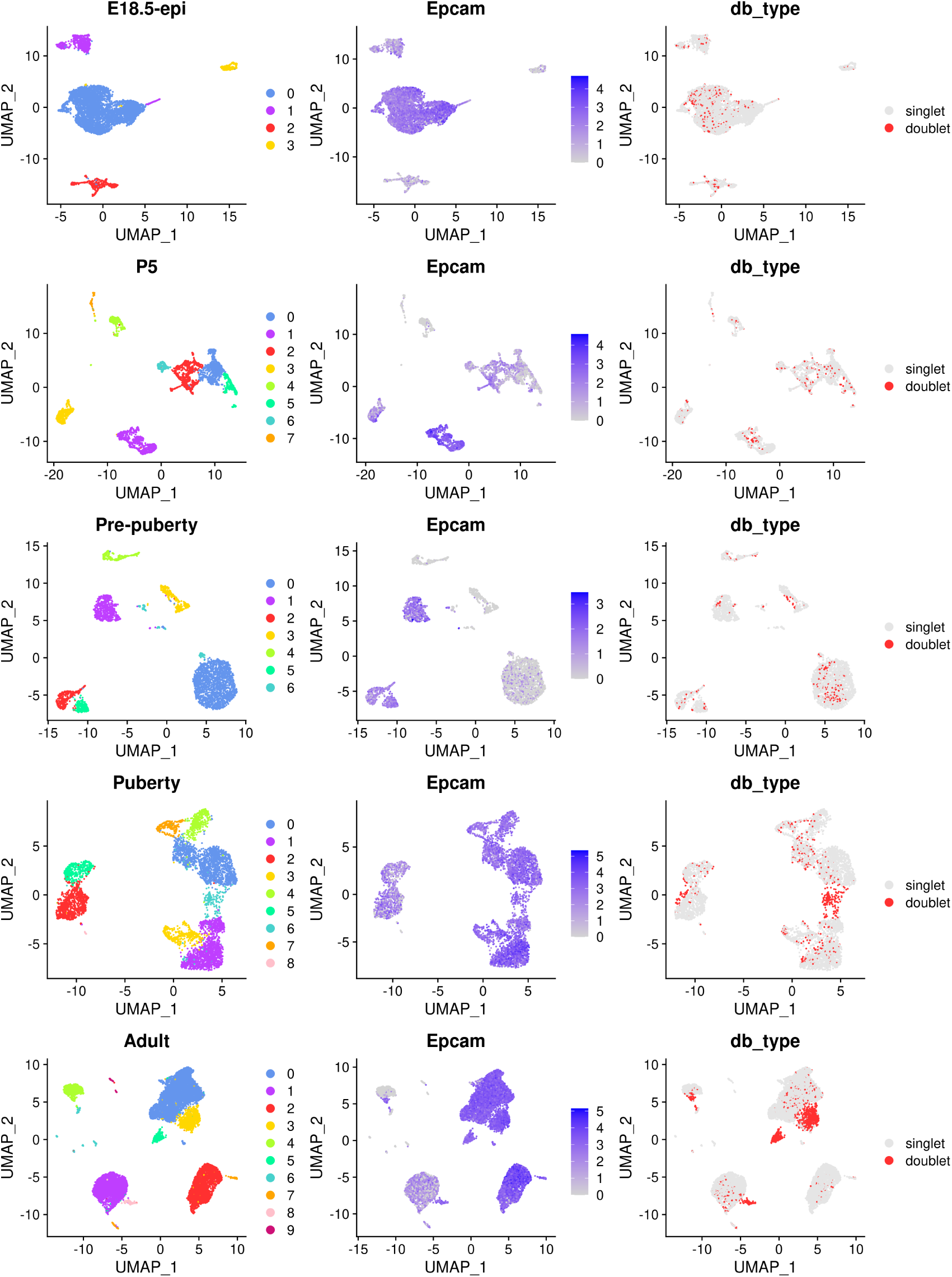
UMAP visualization of each individual samples. The UMAP plots, in sequence from the top row to the bottom row, correspond to E18.5-epi, P5, Pre-puberty, Puberty, and Adult, respectively. In each row, cells are coloured by cluster on the left, by Epcam expression level in the middle, and by doublet prediction on the right.

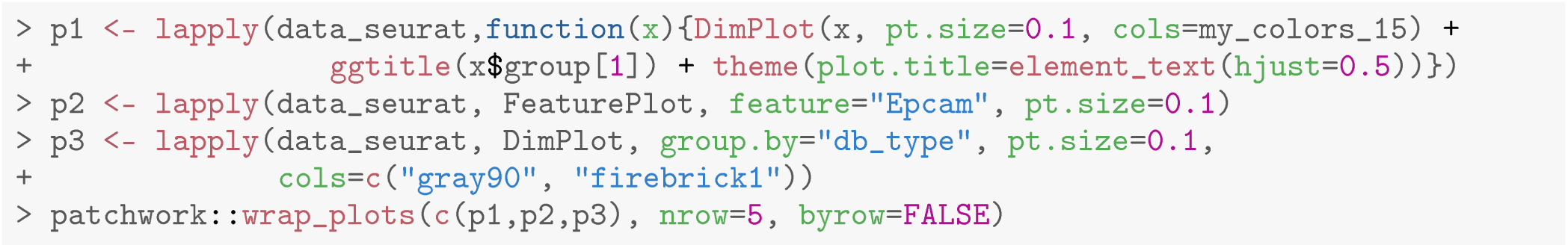

By examining the expression level of the *Epcam* gene, we select the following clusters in each sample as the epithelial cell population.

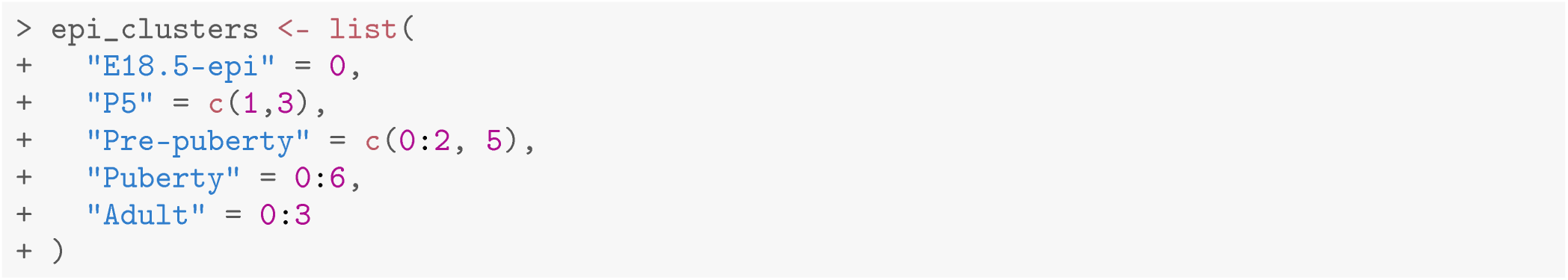

Cells that are non-epithelial and those identified as potential doublets by *scDblFinder* are excluded from the subsequent analysis. The cellular barcodes of the remaining epithelial cells from each sample are stored in the list object called `epi_cells`. The respective number of epithelial cells that are retained for each sample is shown below.

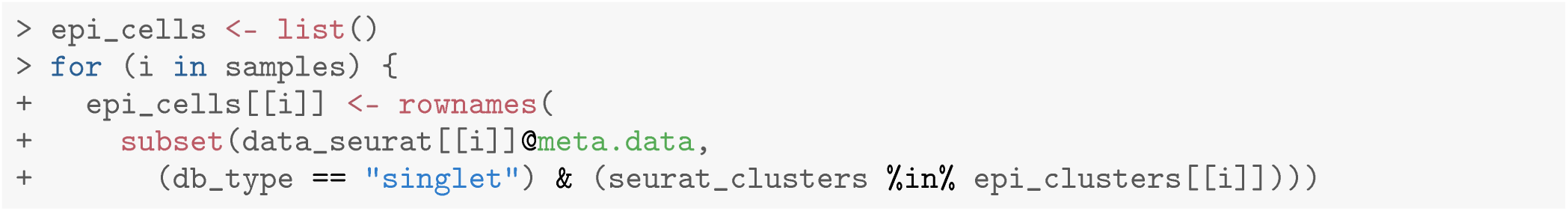

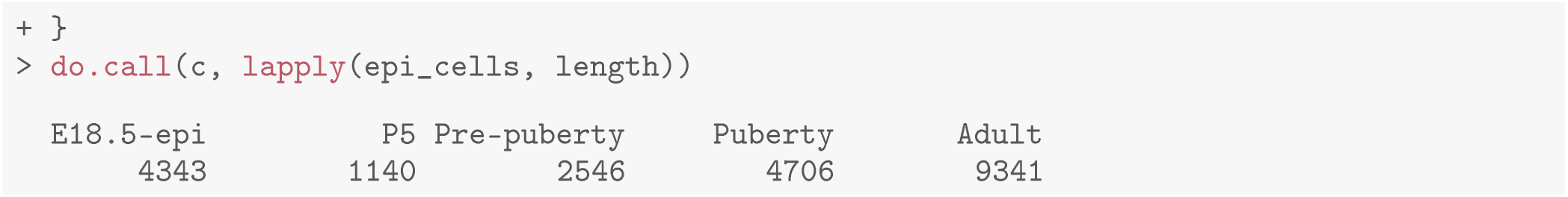

## Data integration

### Integrating epithelial cells of five samples

Since we have five individual scRNA-seq samples, conducting an integration analysis is necessary to explore all cells across these samples simultaneously. In this workflow, we use the anchor-based method in the *Seurat* package for integration. A `Seurat` object is first created from the merged `DGEList` object of epithelial cells using `CreateSeuratObject` function without filtering any cells (`min.features` is set to 0).

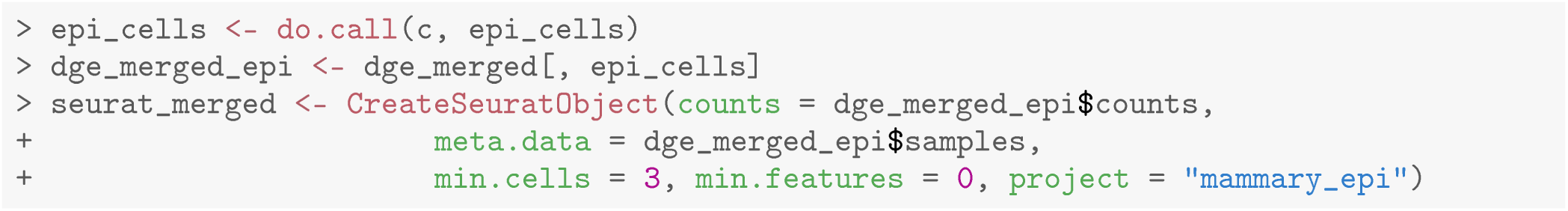

Then the `Seurat` object is split into a list of five `Seurat` objects, where each object corresponds to one of the five samples. For each sample, the log normalization method is applied to normalize the raw count by `NormalizeData`, and highly variable genes are identified by `FindVariableFeatures`.

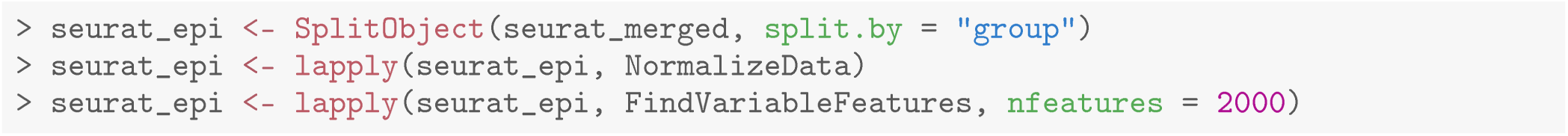

The feature genes used for integration are chosen by `SelectintegrationFeatures`, and these genes are used to identify anchors for integration by `FindintegrationAnchors`. The integration process is performed by `integrateData` based on the identified anchors.

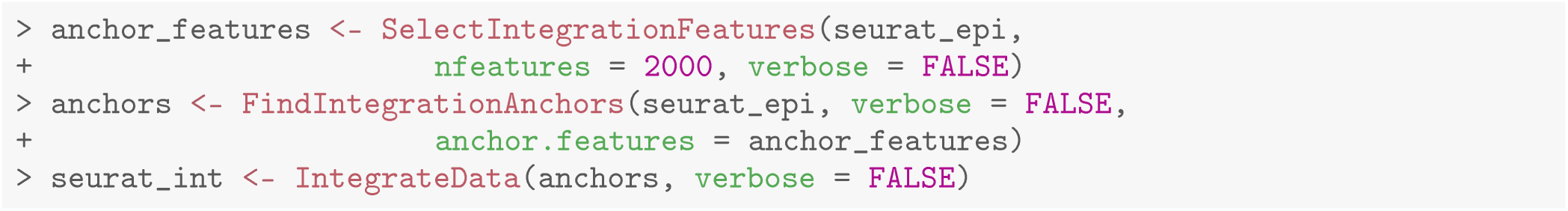

The integrated data are then scaled to have a mean of 0 and a variance of 1 by `ScaleData`. PCA is performed on the scaled data using `RunPCA`, followed by UMAP using `RunUMAP`. Cell clusters of the integrated data are identified by using `FindNeighbors` and `FindClusters`.

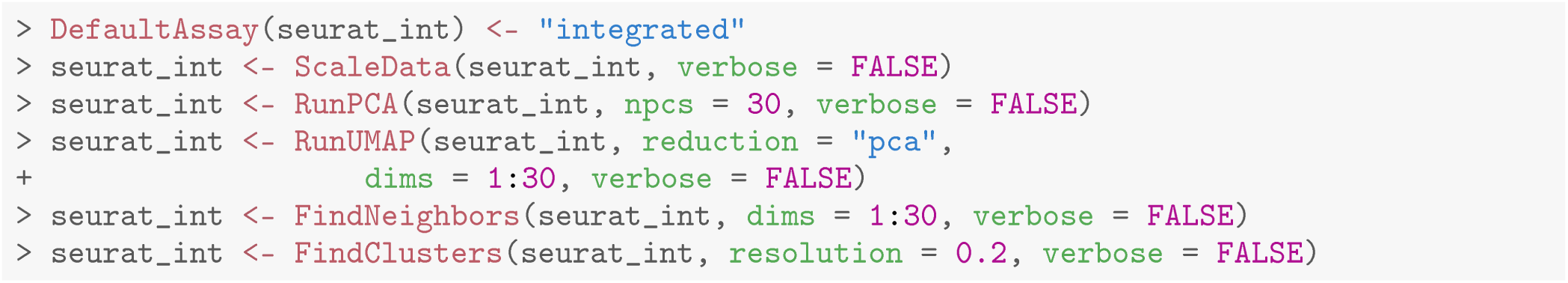

UMAP plots are generated to visualize the integration and cell clustering results (Figure 3). The UMAP plot indicates the presence of three major cell clusters (cluster 0, 1, and 2), which are bridged by intermediate clusters located in between them. Cells at the later stages largely dominate the three major cell clusters, while cells at the earlier stages are predominantly present in the intermediate clusters in the middle.

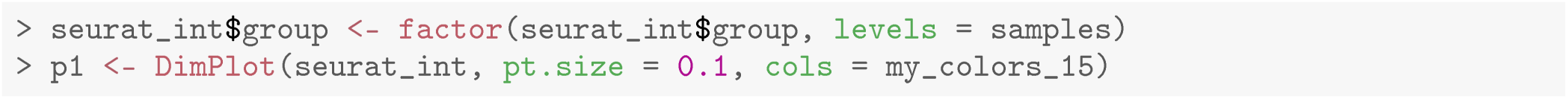

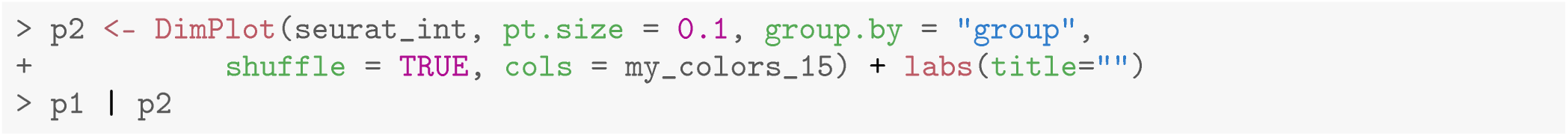

**Figure 3.**
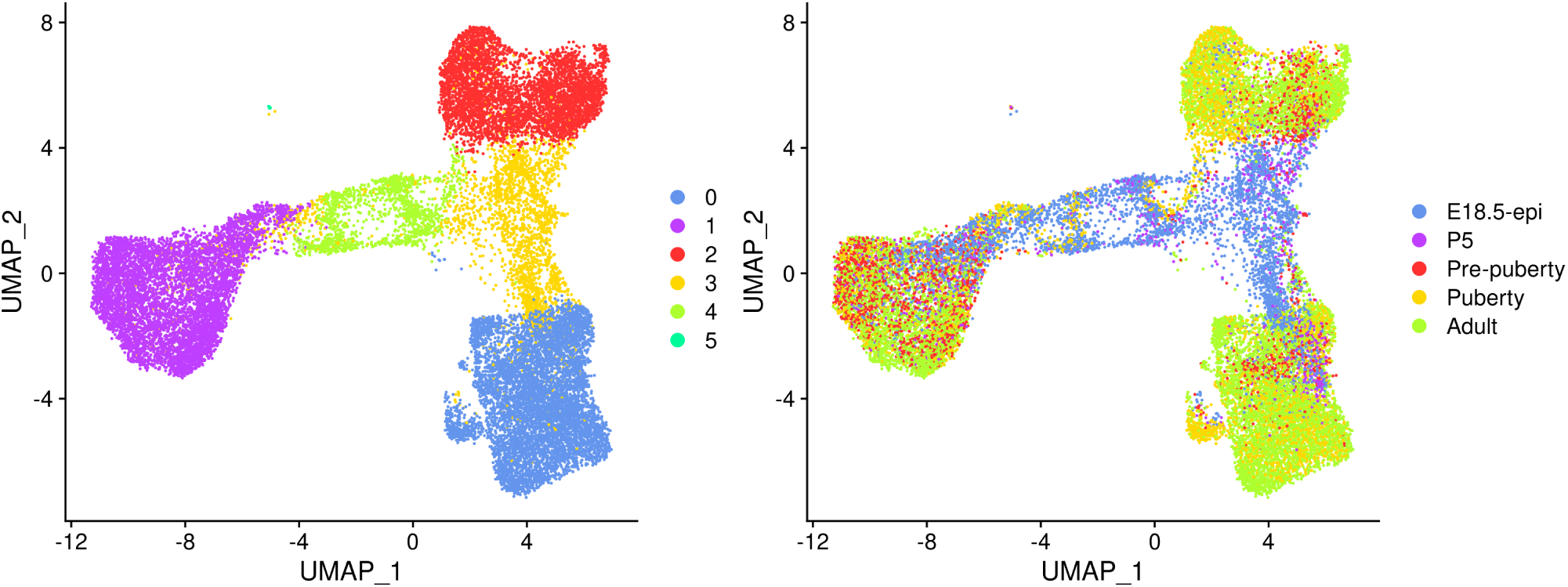
UMAP visualization of the integrated data. Cells are coloured by cluster on the left and by original sample on the right.

### Cell type identification

The mammary gland epithelium consists of three major cell types: basal myoepithelial cells, luminal progenitor (LP) cells and mature luminal (ML) cells. These three major epithelial cell populations have been well studied in the literature. By examining the classic marker genes of the three cell types, we are able to identify basal, LP and ML cell populations in the integrated data (Figure 4). Here we use *Krt14* and *Acta2* for basal, *Csn3* and *Elf5* for LP, and *Prlr* and *Areg* for ML. We also examine the expression level of *Hmgb2* and *Mki67* as they are typical markers for cycling cells and the expression level of *Igfbp7* and *Fabp4* as they are marker genes for stromal cells.

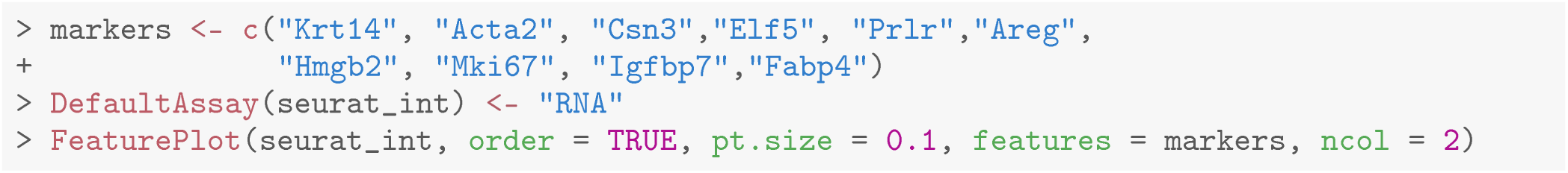

**Figure 4.**
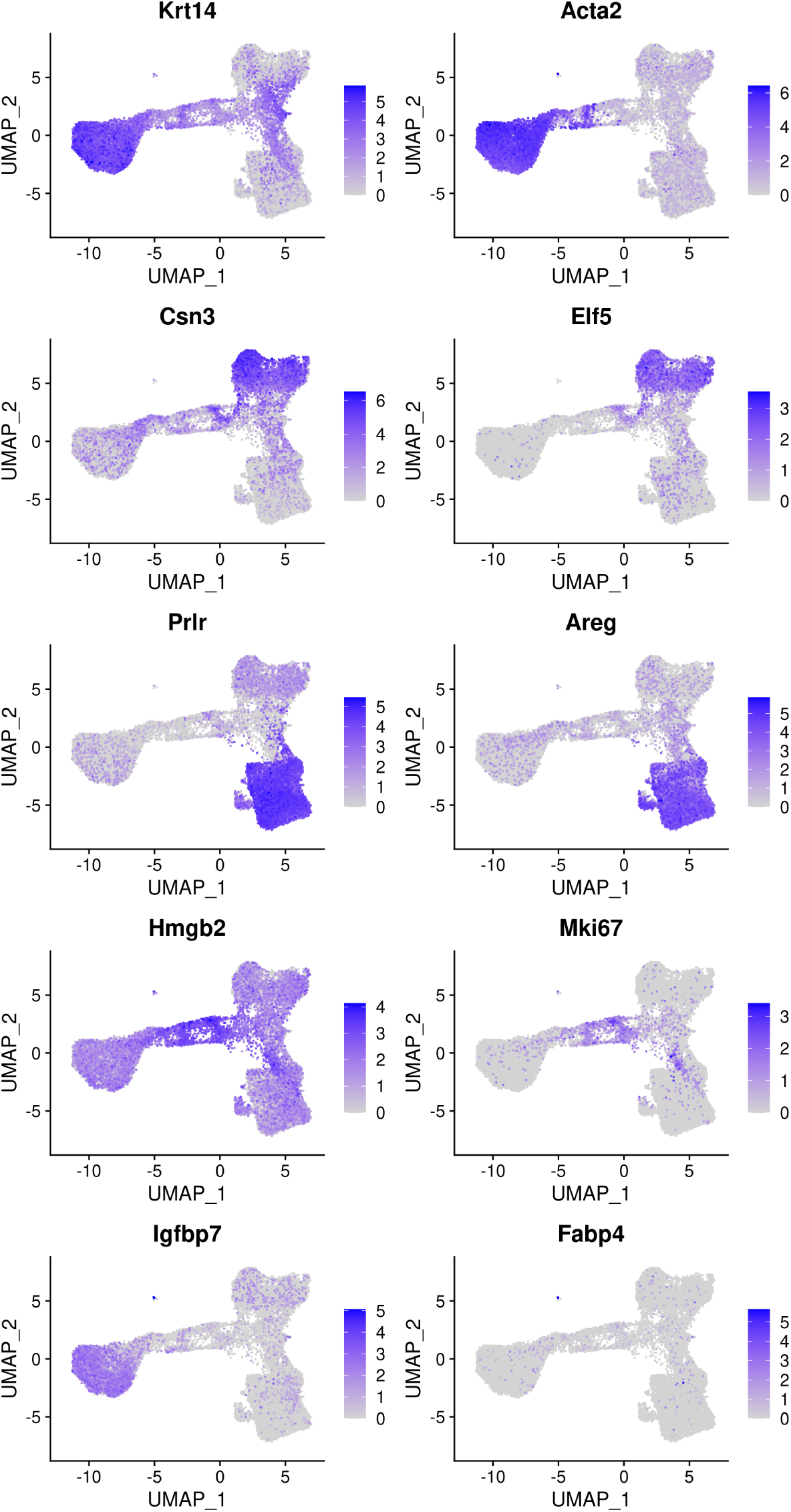
Feature plots of the integrated data. Genes from the top row to the bottom rows are the markers of basal, LP, ML, cycling cells, and stroma, respectively.

Based on the feature plots, cluster 1, cluster 2 and cluster 0 represent the basal, LP and ML cell populations, respectively. Cluster 4 mainly consists of cycling cells, whereas cluster 3 seems to be a luminal intermediate cell cluster expressing both LP and ML markers. Cluster 5 consists of a few non-epithelial (stromal) cells that have not been filtered out previously.

The number of cells in each cluster for each sample is shown below.

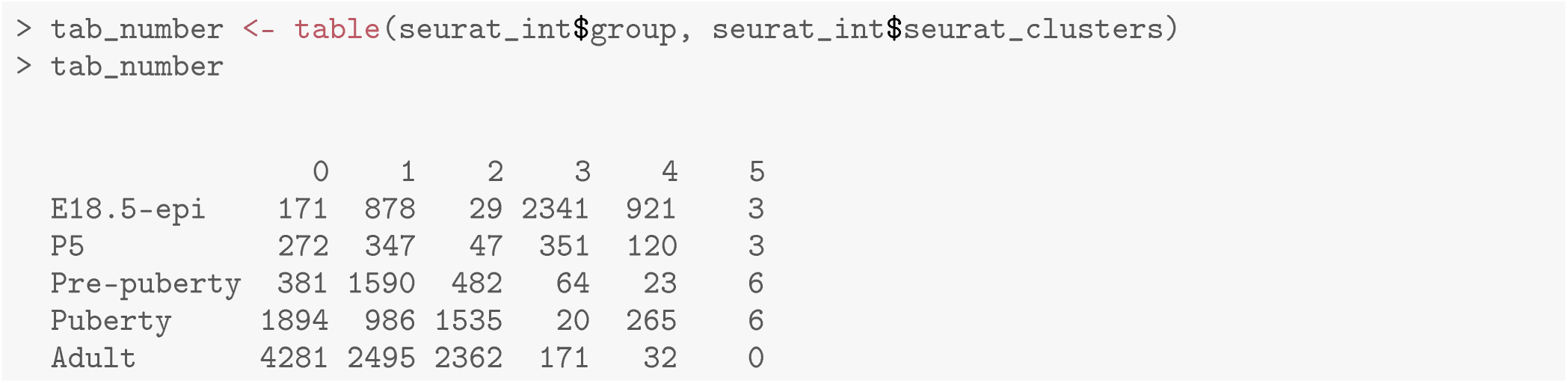

The proportion of cells in each cluster is calculated for each sample to compare the variation in cell composition across different stages.

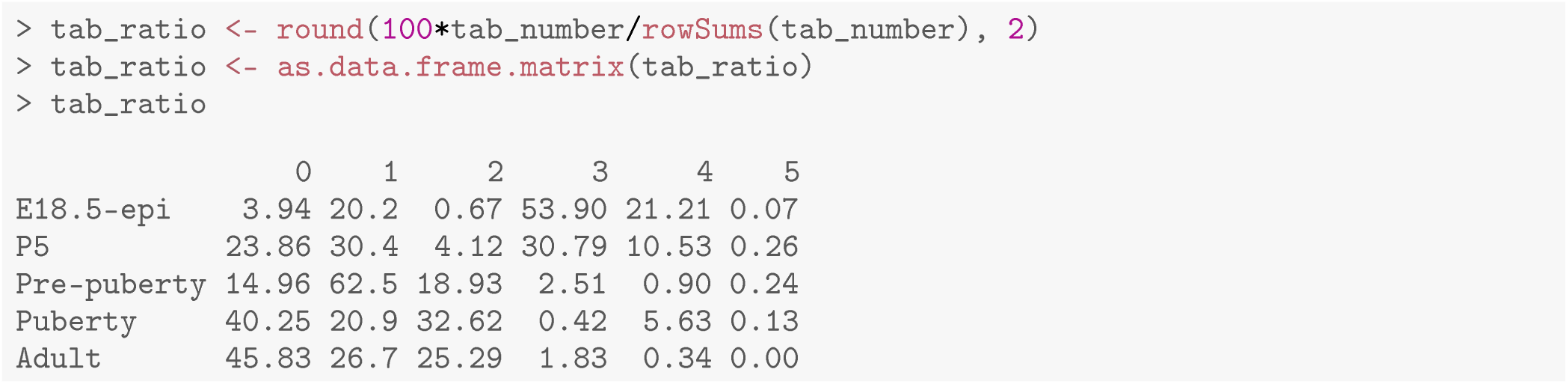

The bar plot (Figure 5) shows the proportion of different cell types in samples at different developmental stages. Specifically, the proportion of basal cells (purple) demonstrates an ascending trend from E18.5 to pre-puberty stage, after which it declines towards adult stage. The LP cell proportion (red) rises from E18.5 to puberty stage, followed by a slight dip at adult stage. Although the proportion of ML cells (blue) is higher at P5 than pre-puberty stage, it shows an overall increasing trend. Cycling cells (green) constitute the highest proportion at E18.5 stage, but decrease to a smaller proportion at pre-puberty stage, with a slight increase at puberty stage, and subsequently, they reduce to a negligible proportion at adult stage. The augmented cycling cell proportion at puberty stage aligns with the ductal morphogenesis characteristics of the mammary gland. The luminal intermediate cell proportion (yellow) displays a decreasing trend from E18.5 stage to adult stage.

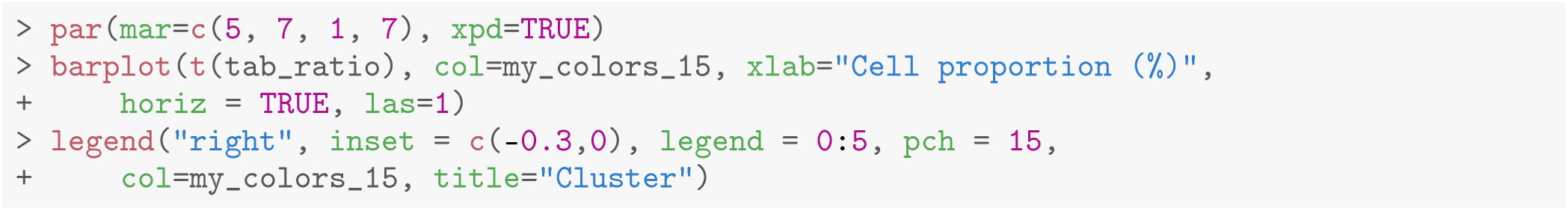

**Figure 5.**
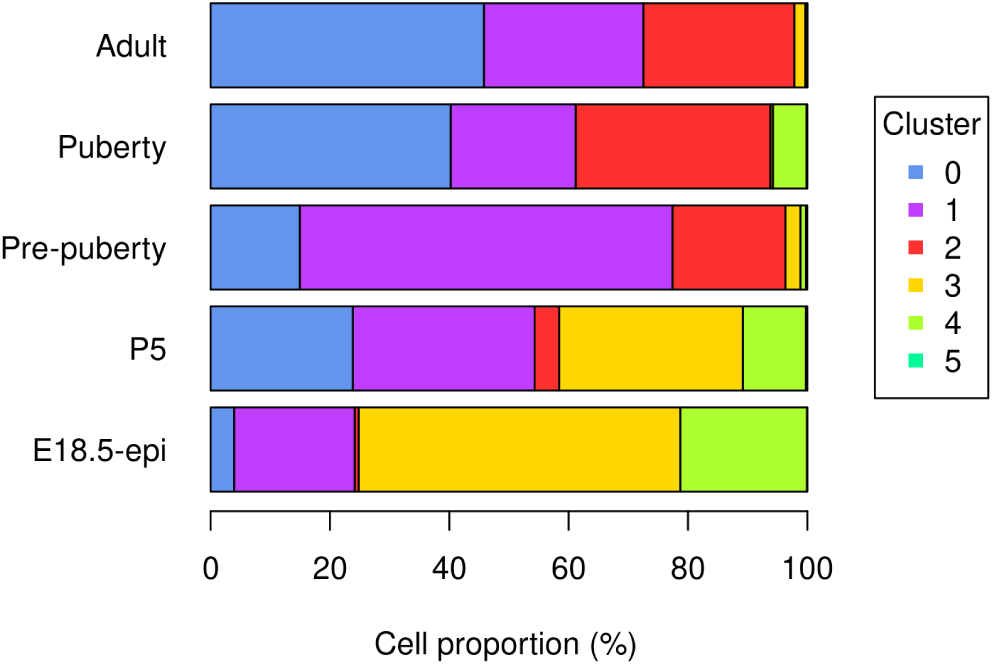
Bar plot of cell proportion of each cluster in each sample.

## Trajectory analysis with monocle3

### Constructing trajectories and pseudotime

Many biological processes manifest as a dynamic sequence of alterations in the cellular state, which can be estimated through a “trajectory” analysis. Such analysis is instrumental in detecting the shifts between different cell identities and modeling gene expression dynamics. By treating single-cell data as a snapshot of an uninterrupted process, the analysis establishes the sequence of cellular states that forms the process trajectory. The arrangement of cells along these trajectories can be interpreted as pseudotime.

Here, we use the *monocle3* package to infer the development trajectory in the mouse mammary gland epithelial cell population. The Seurat object of the integrated data is first converted into a `cell_data_set` object to be used in *monocle3*.

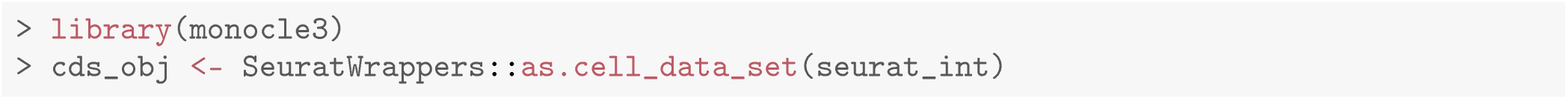

*monocle3* re-clusters cells to assign them to specific clusters and partitions, which are subsequently leveraged to construct trajectories. If multiple partitions are used, each partition will represent a distinct trajectory. The calculation of pseudotime, which indicates the distance between a cell and the starting cell in a trajectory, is conducted during the trajectory learning process. These are done using the `cluster_cells` and `learn_graph` functions. To obtain a single trajectory and avoid a loop structure, both `use_partition` and `close_loop` are turned off in `learn_graph`.

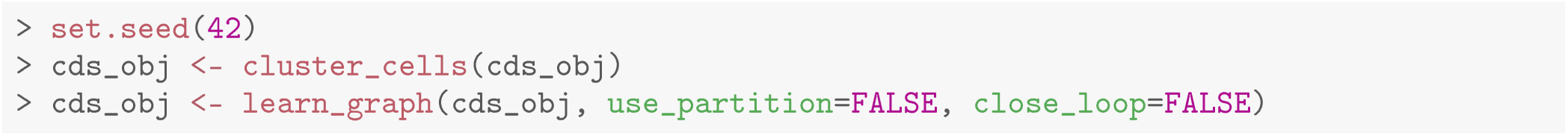

### Visualizing trajectories and pseudotime

The `plot_cells` function of *monocle3* is used to generate a trajectory plot that superimposes the trajectory information onto the UMAP representation of the integrated data. By adjusting the `label_principal_points` parameter, the names of roots, leaves, and branch points can be displayed. Cells in the trajectory UMAP plot (Figure 6) on the left are colored by cell cluster identified in the previous Seurat integration analysis.

**Figure 6.**
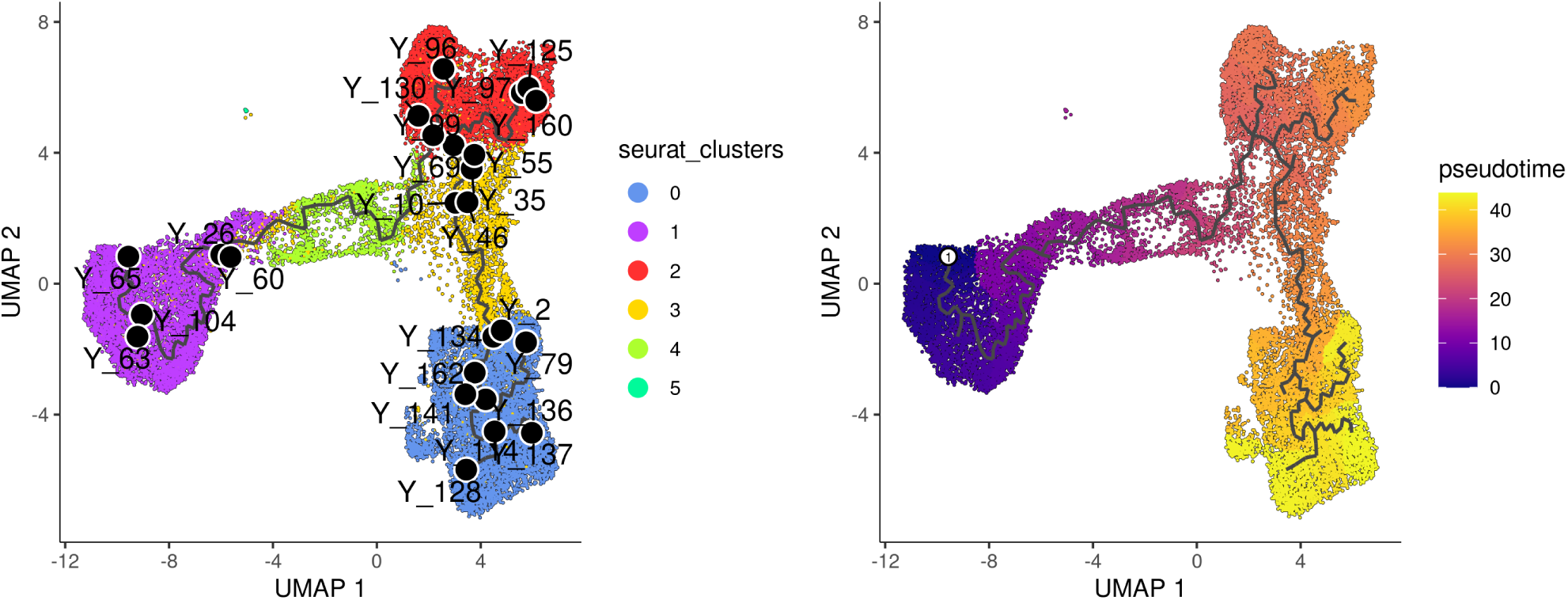
UMAP visualization of trajectory and pseudotime computed by monocle3. Cells are coloured by cluster on the left and by pseudotime on the right.

Along the *monocle3* trajectory analysis, several nodes are identified and marked with black circular dots on the resulting plot, representing key points along the trajectories. To establish the order of cells and calculate their corresponding pseudotime, it is necessary to select a starting node from among the identified nodes. For this analysis, node `IIY_65II` in the basal population (cluster 1) was selected as the starting node, as mammary stem cells are known to be enriched in the basal population and give rise to LP and ML cells in the epithelial lineage [19]. It should be noted that node numbers may vary depending on the version of monocle3 used.

The cells are then ordered and assigned pseudotime values by the `order_cells` function in *monocle3*. The resulting pseudotime information can be visualized on the UMAP plot by using the `plot_cells` function, as demonstrated in the UMAP plot on the right (Figure 6).

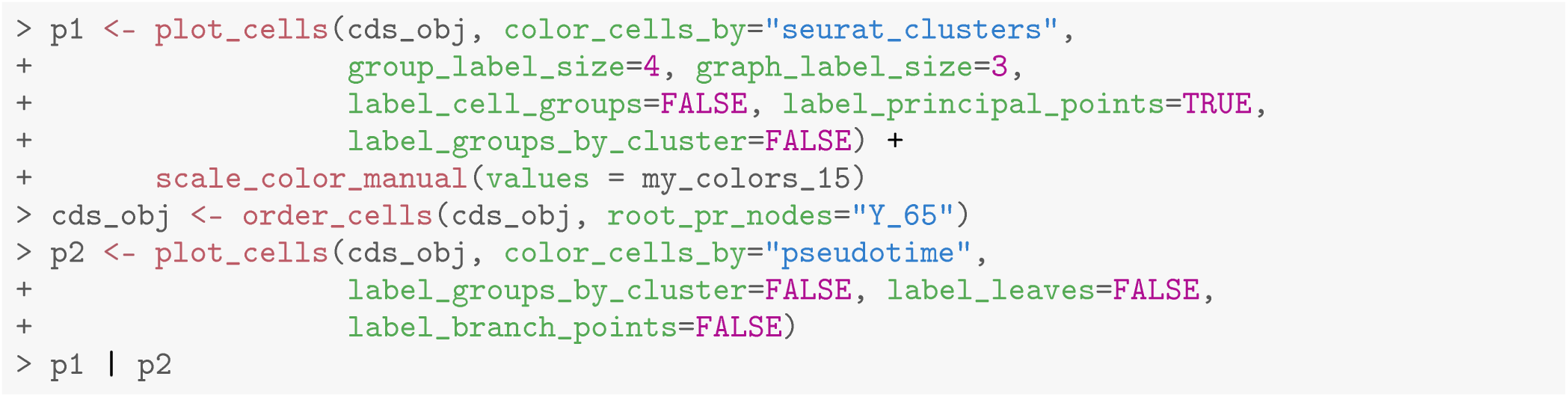

The `pseudotime` function in *monocle3* allows users to extract the pseudotime values of the cells from a `cell_data_set` object. This information can then be stored in the metadata of the Seurat object for further analysis.

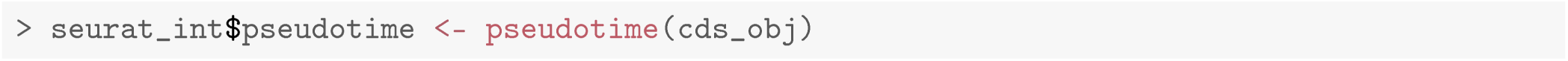

## Psuedo-bulk time course analysis with edgeR

### Constructing pseudo-bulk profiles

After obtaining the pseudotime of each cell, we proceed to a time course analysis to identify genes that change significantly along the pseudotime time. Our approach involves creating pseudo-bulk samples using a pseudo-bulking approach and performing an *edgeR*-style time course analysis.

To create the pseudo-bulk samples, read counts are aggregated for all cells with the same combination of sample and cluster. The number of cells used to construct each pseudo-bulk sample is added to the sample metadata. The average pseudotime of all cells in each pseudo-bulk sample is used as the pseudotime for that sample.

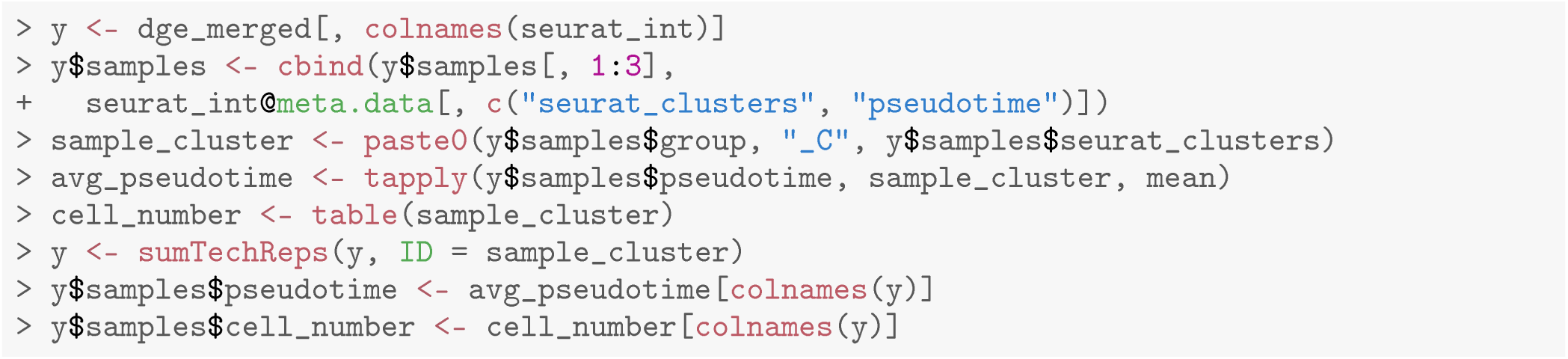

The Entrez gene IDs are added to the gene information. Genes with no valid Entrez gene IDs are removed from the downstream analysis.

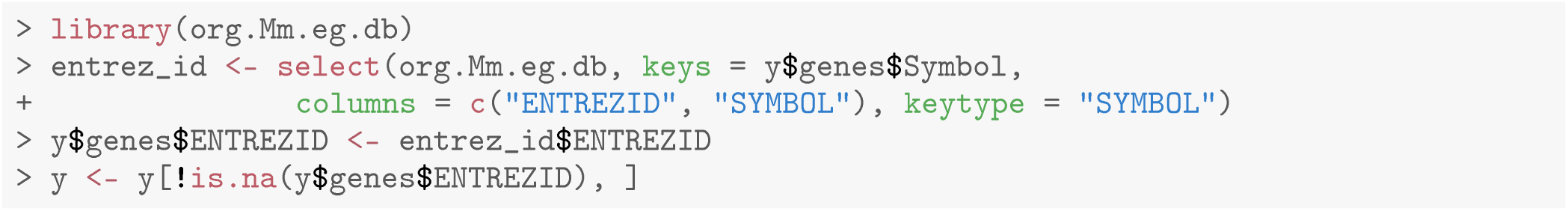

The samples are ordered by average pseudotime for the following analysis.

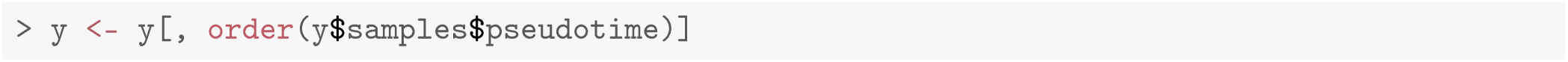

### Filtering and normalization

We now proceed to the standard *edgeR* analysis pipeline, which starts with filtering and normalization. The sample information, such as library sizes, average pseudotime and cell numbers, are shown below.

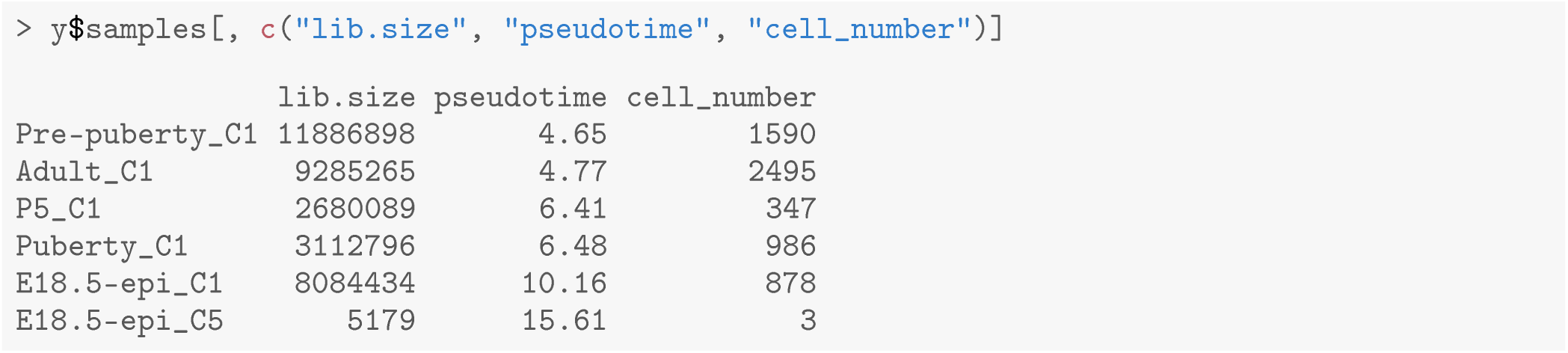

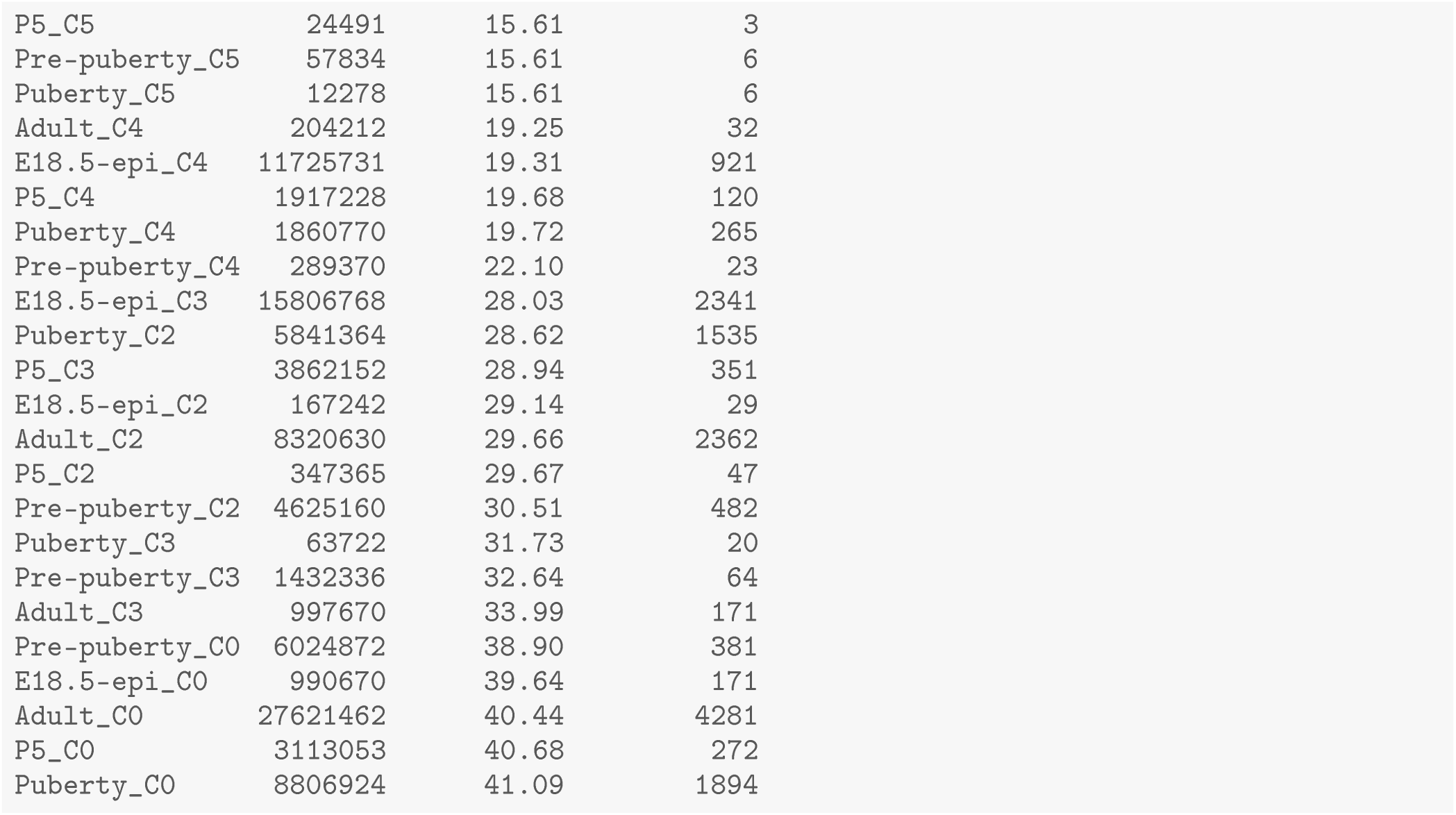

To ensure the reliability of the analysis, it is recommended to remove pseudo-bulk samples that are constructed from a small number of cells. We suggest each pseudo-bulk sample should contain at least 30 cells. In this analysis, we identified seven pseudo-bulk samples that were constructed with less than 30 cells and removed them form the analysis.

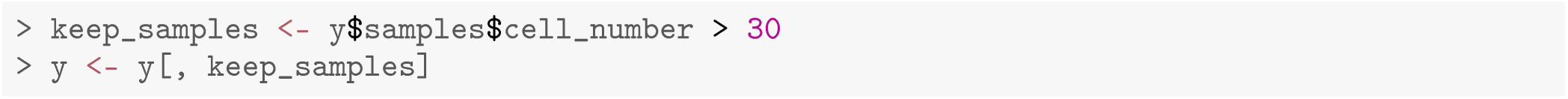

Genes with very low count number are also removed from the analysis. This is performed by the `filterByExpr` function in *edgeR*.

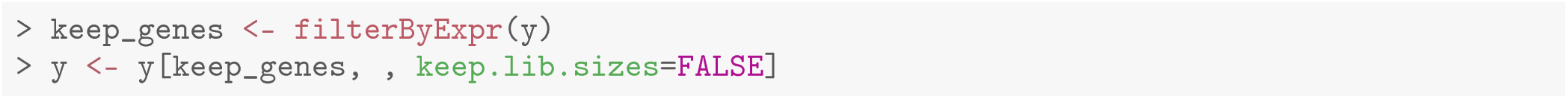

The number of genes and samples after filtering are shown below.

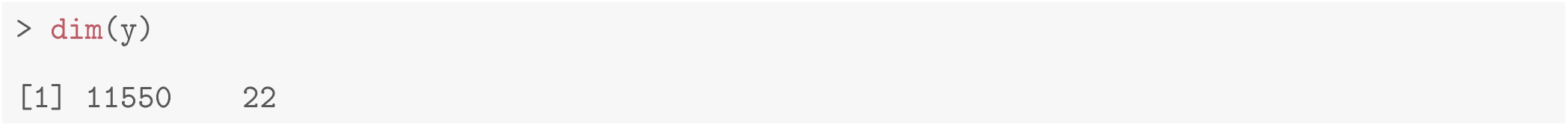

Normalization is performed by the trimmed mean of M values (TMM) method [20] implemented in the `calcNormFactors` function in *edgeR*.

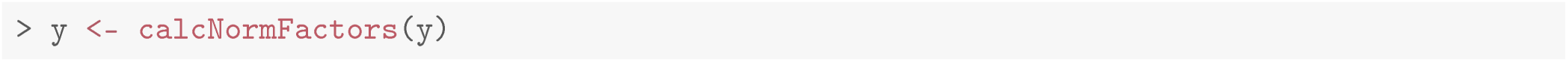

A Multi-dimensional scaling (MDS) plot serves as a valuable diagnostic tool for investigating the relationship among samples. MDS plots are produced using the `plotMDS` function in *edgeR* (Figure 7).

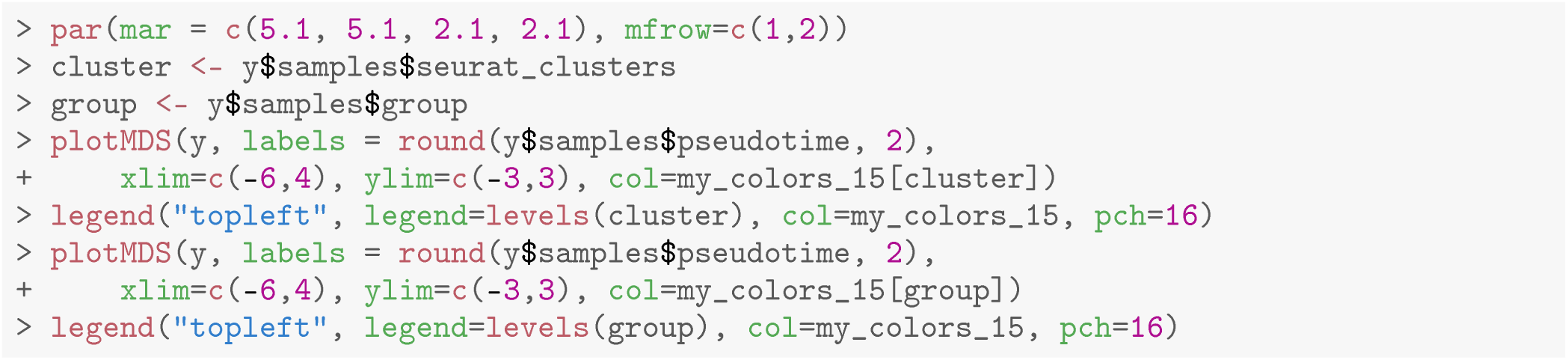

**Figure 7.**
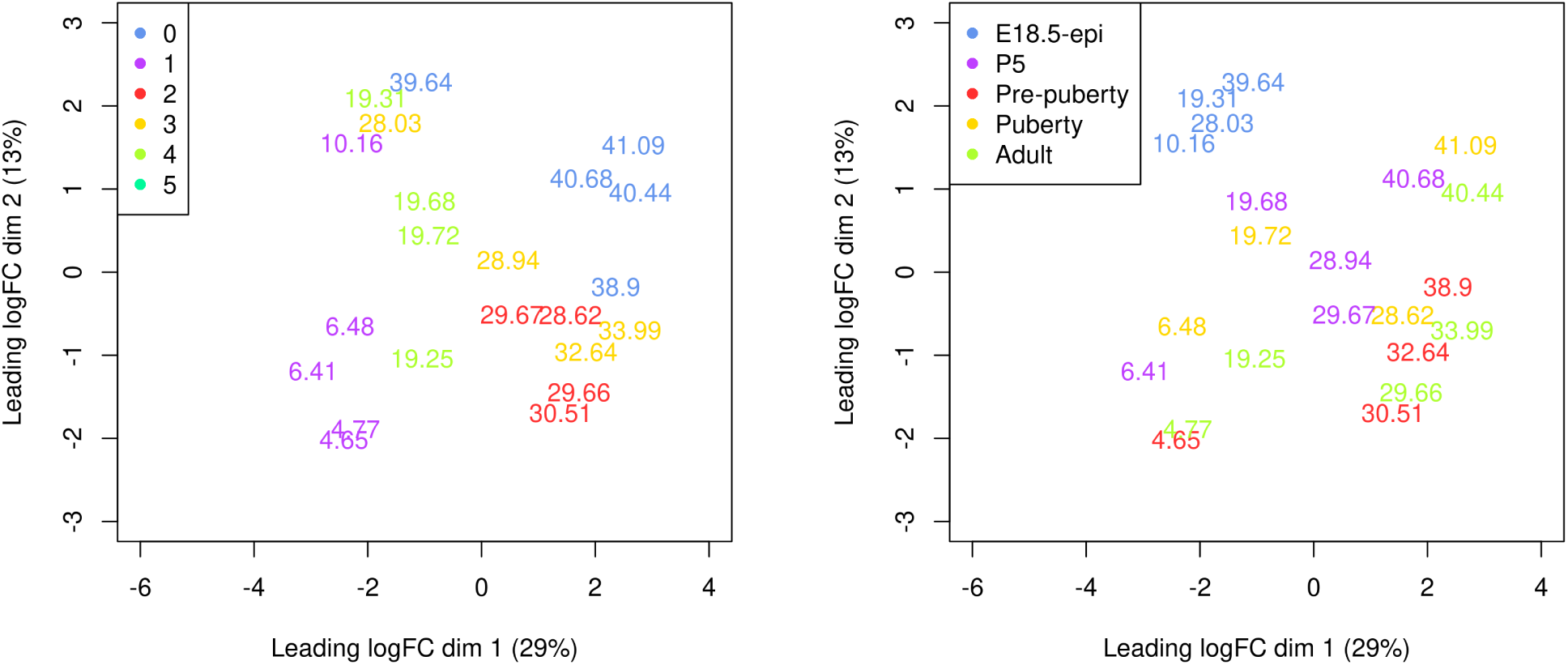
Multi-dimensional scaling (MDS) plot of the pseudo-bulk samples labelled by pseudotime. Samples are coloured by origianl cell cluster on the left and by developmental stage on the right.

On the MDS plot, pseudo-bulk samples derived from the same cell cluster are close to each other. The samples are positioned in ascending order of pseudotime from left to right, suggesting a continuous shift in the gene expression profile throughout the pseudotime.

### Design matrix

The aim of a time course experiment is to examine the relationship between gene abundances and time points. Assuming gene expression changes smoothly over time, we use a natural cubic spline with degrees of freedom of 3 to model gene expression along the pseudotime. The spline design matrix is generated by `ns` function in *splines*. The design matrix is also reformed so that the first column represents the linear trend.

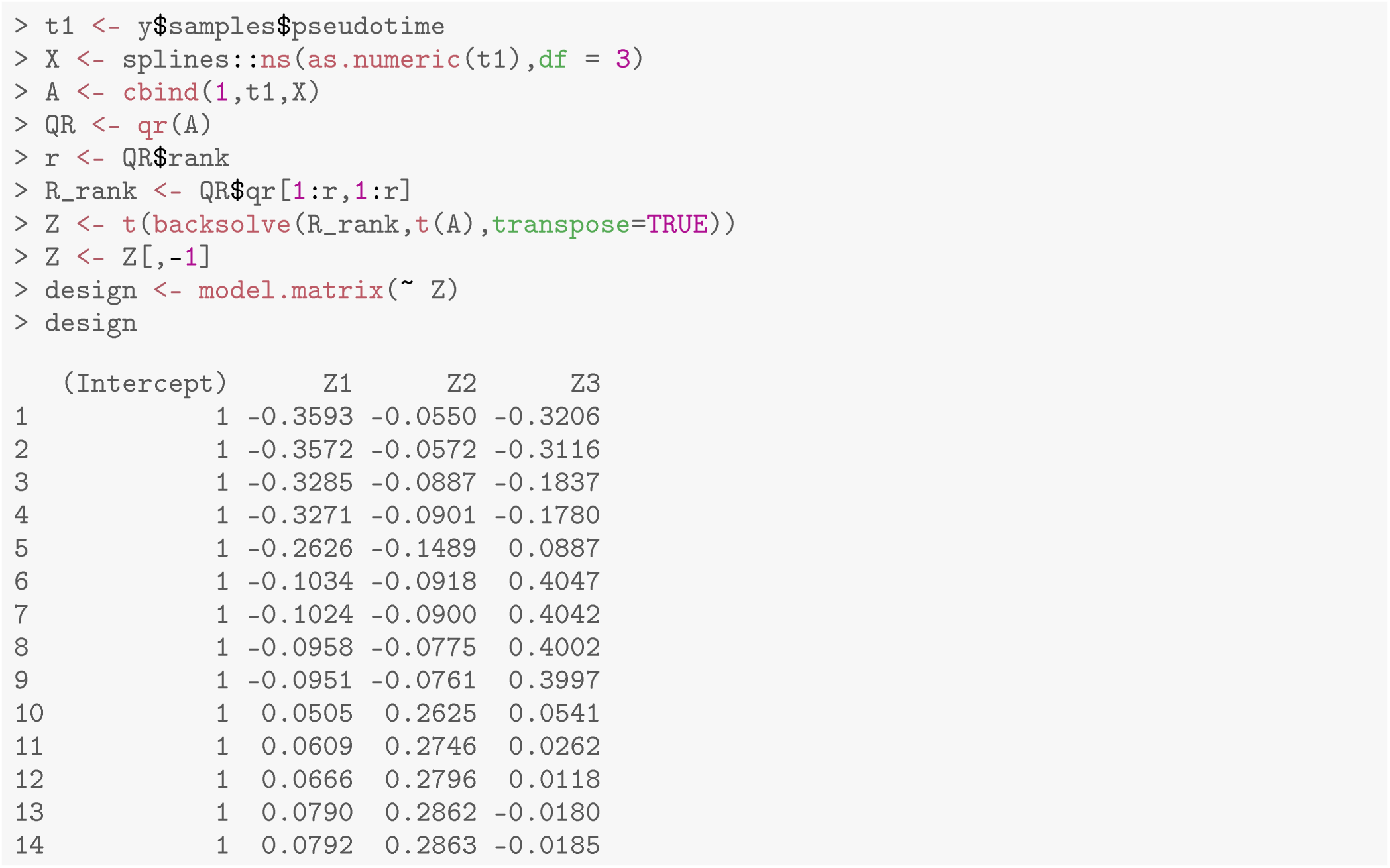

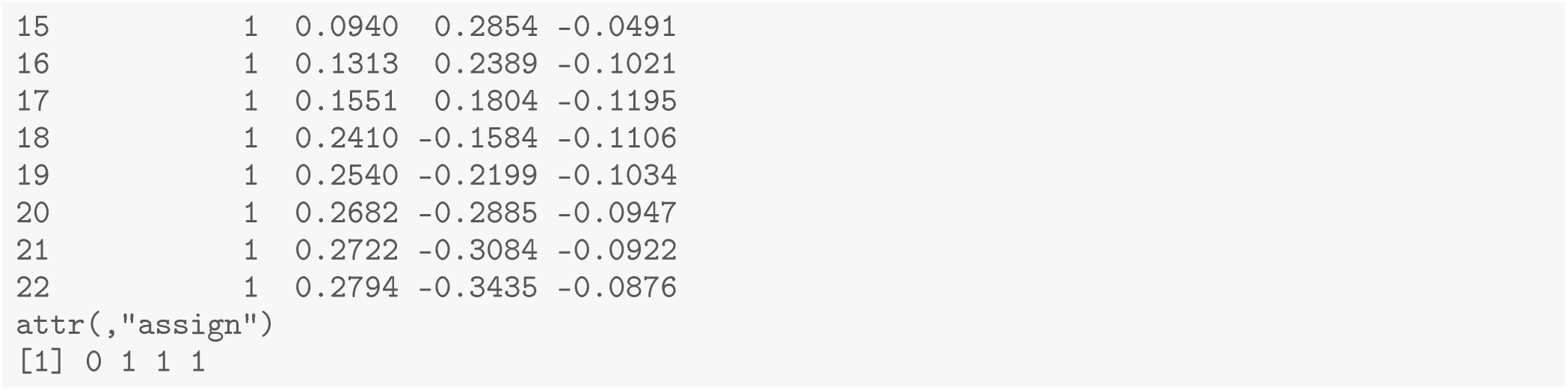

### Dispersion estimation

The *edgeR* package uses negative binomial (NB) distribution to model read counts of each gene across all the sample. The NB dispersions are estimated by the `estimateDisp` function. The estimated common, trended and gene-specific dispersions can be visualized by `plotBCV` (Figure 8).

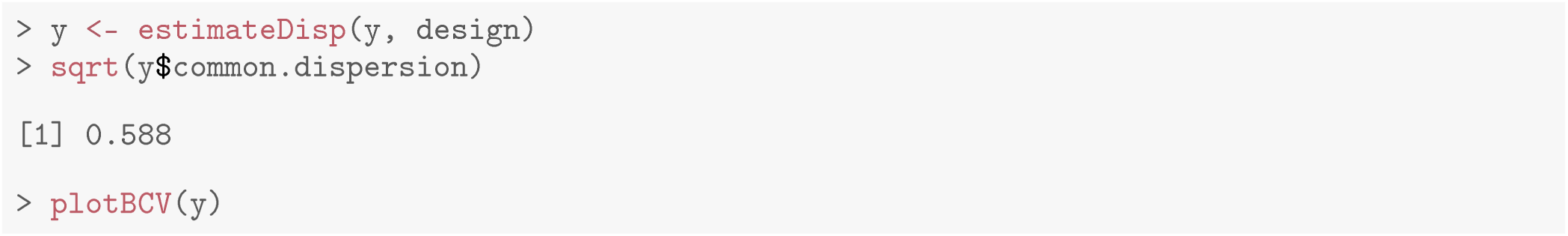

**Figure 8.**
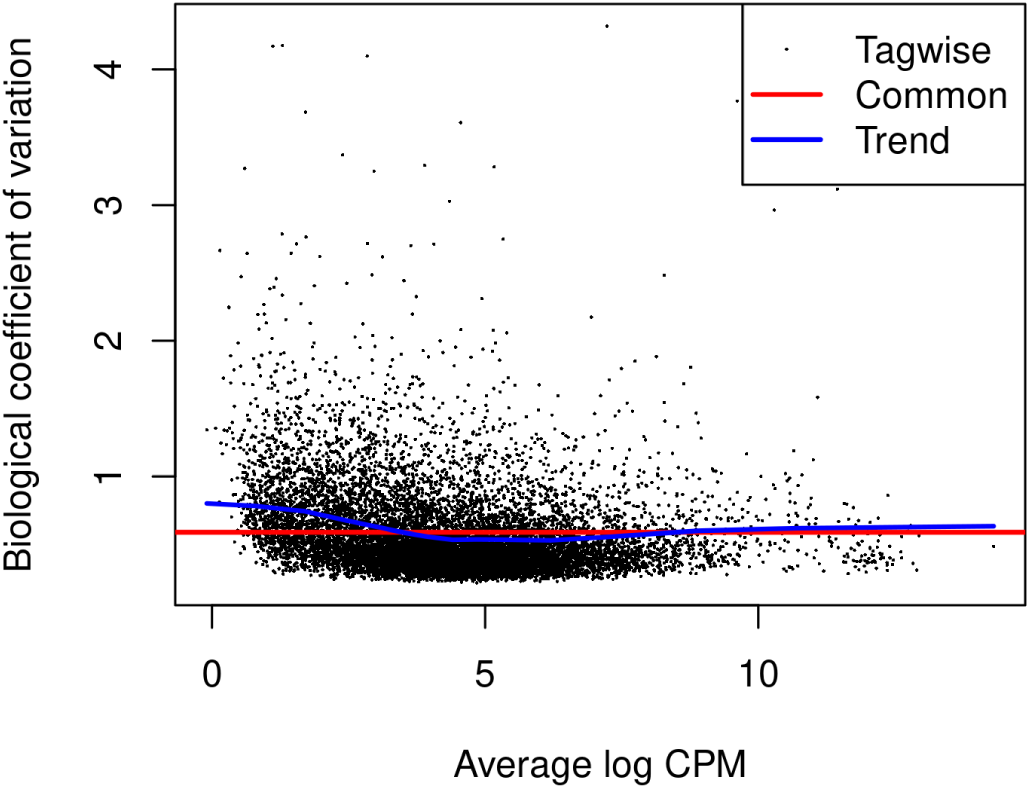
A scatter plot of the biological coefficient of variation (BCV) against the average abundance of each gene. The square-root estimates of the common, trended and gene-wise NB dispersions are shown.

The NB model can be extended with quasi-likelihood (QL) methods to account for gene-specific variability from both biological and technical sources [21, 22]. Note that only the trended NB dispersion is used in the QL method. The gene-specific variability is captured by the QL dispersion.

The `glmQLFit` function is used to fit a QL model and estimate QL dispersions. The QL dispersion estimates can be visualized by `plotQLDisp` (Figure 9).

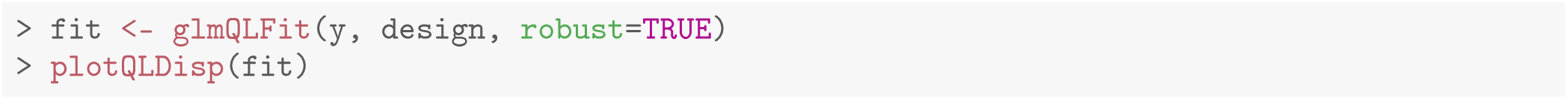

**Figure 9.**
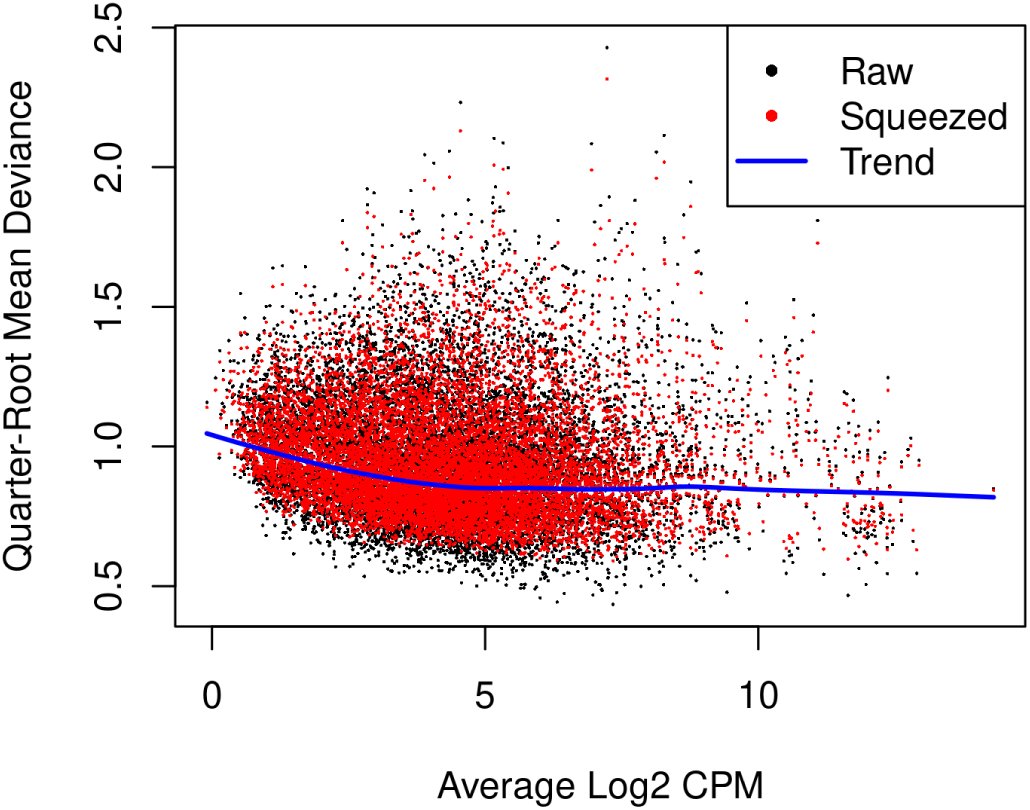
A scatter plot of the quarter-root QL dispersion against the average abundance of each gene. Estimates are shown for the raw, trended and squeezed dispersions.

### Time course trend analysis

The QL F-tests are performed by `glmQLFTest` in *edgeR* to identify genes that change significantly along the pseudotime. The test are conducted on all three covariates of the spline model matrix. This is because the significance of any of the three coefficients would indicate a strong correlation between gene expression and pseudotime.

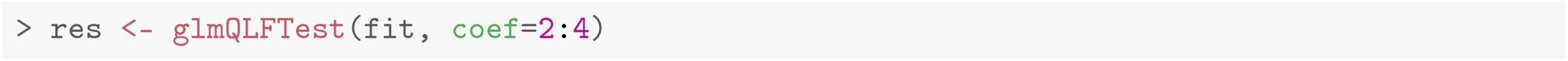

The number of genes significantly associated with pseudotime (FDR < 0.05) are shown below.

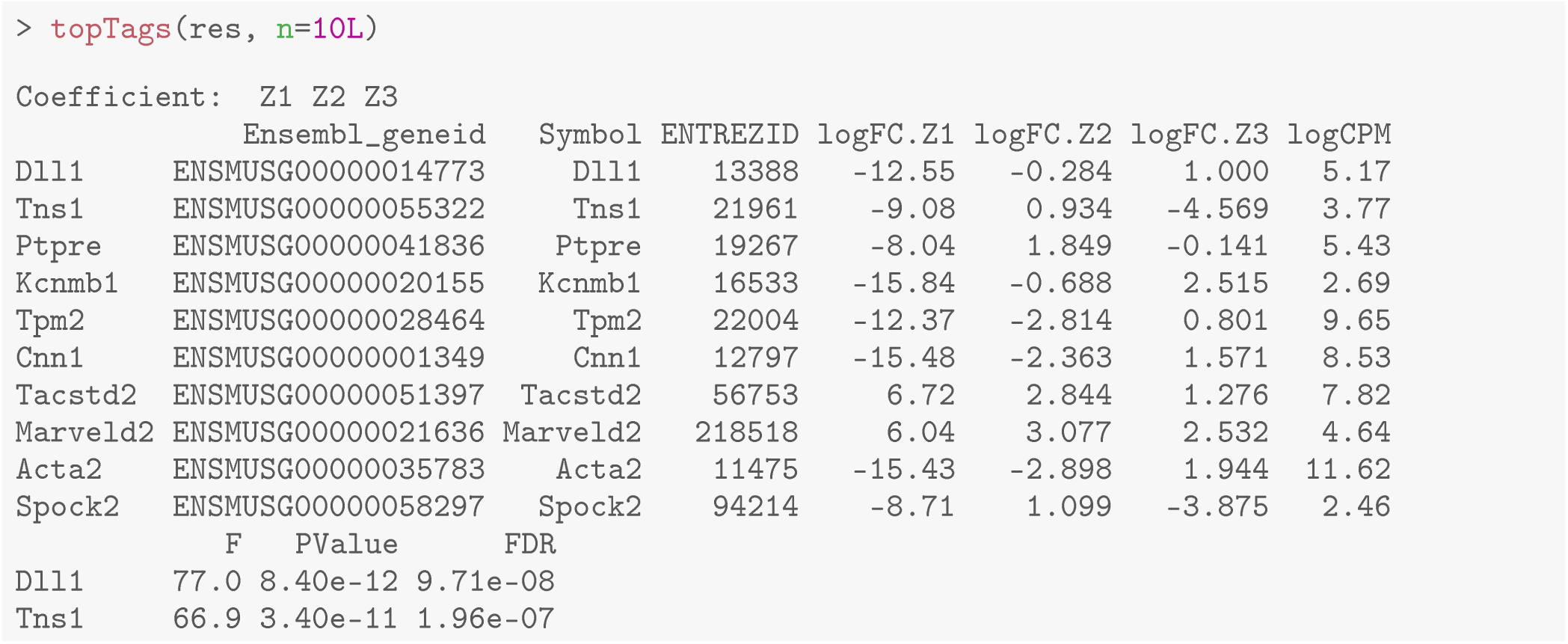

Top significant genes can be viewed by `topTags`.

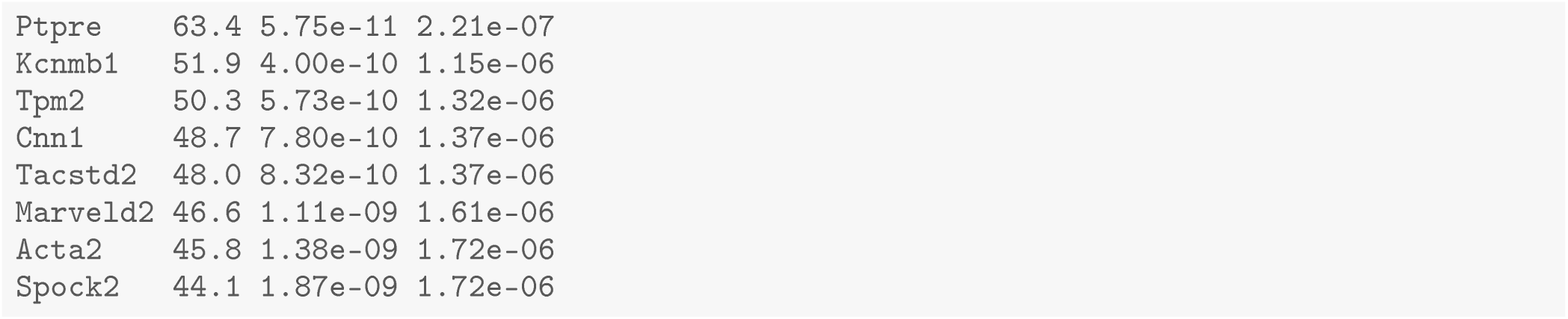

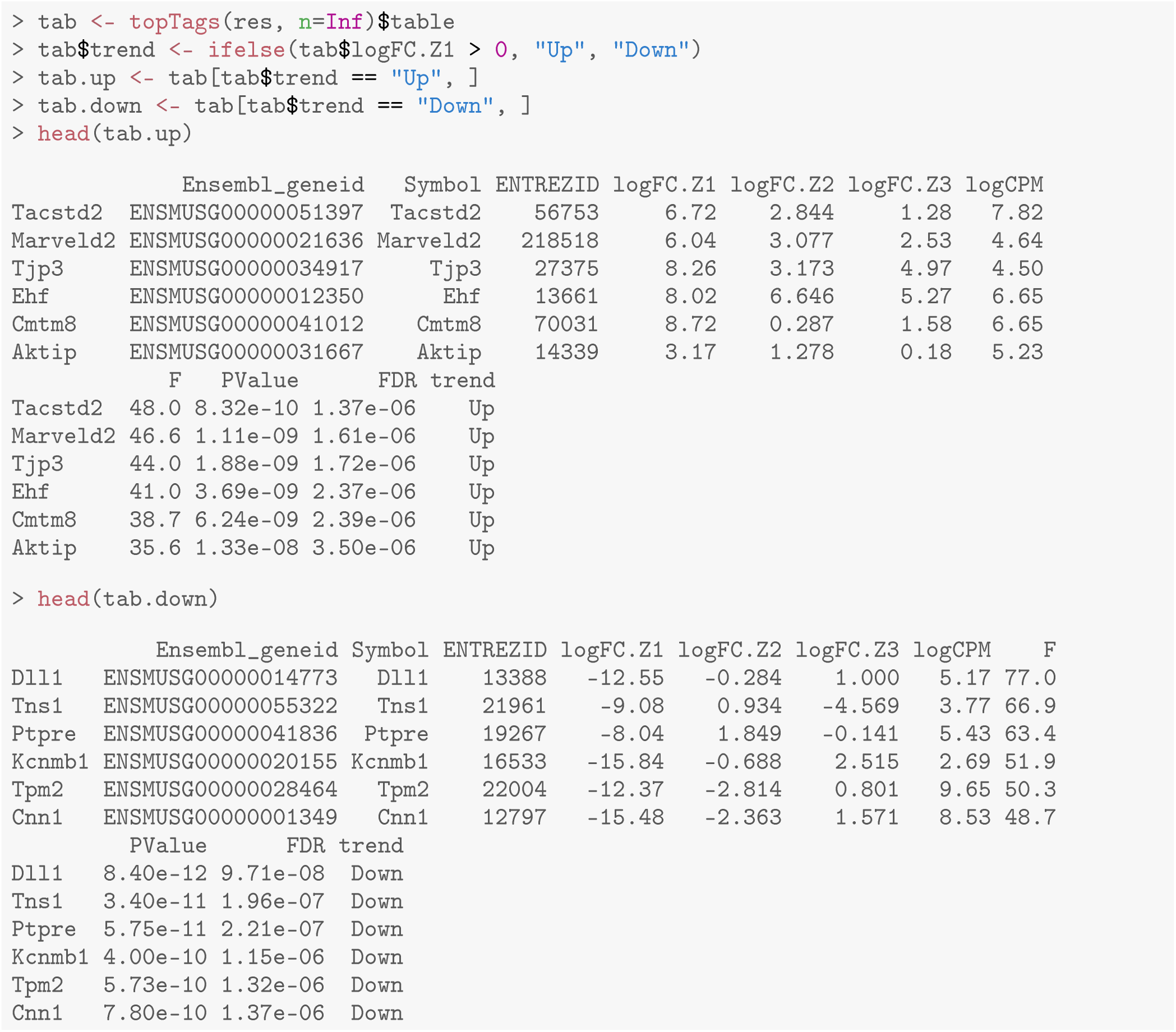

The `logFC.Z1`, `logFC.Z2`, and `logFC.Z3` values in the table above denote the estimated coefficients of Z1, Z2, and Z3 for each gene. It should be noted that these values do not carry the same interpretation as log-fold changes in traditional RNA-seq differential expression analysis. For each gene, the sign of the coefficient `logFC.Z1` indicates whether the expression level of that gene increases or decreases along pseudotime in general. The top increasing and the top decreasing genes are listed below.

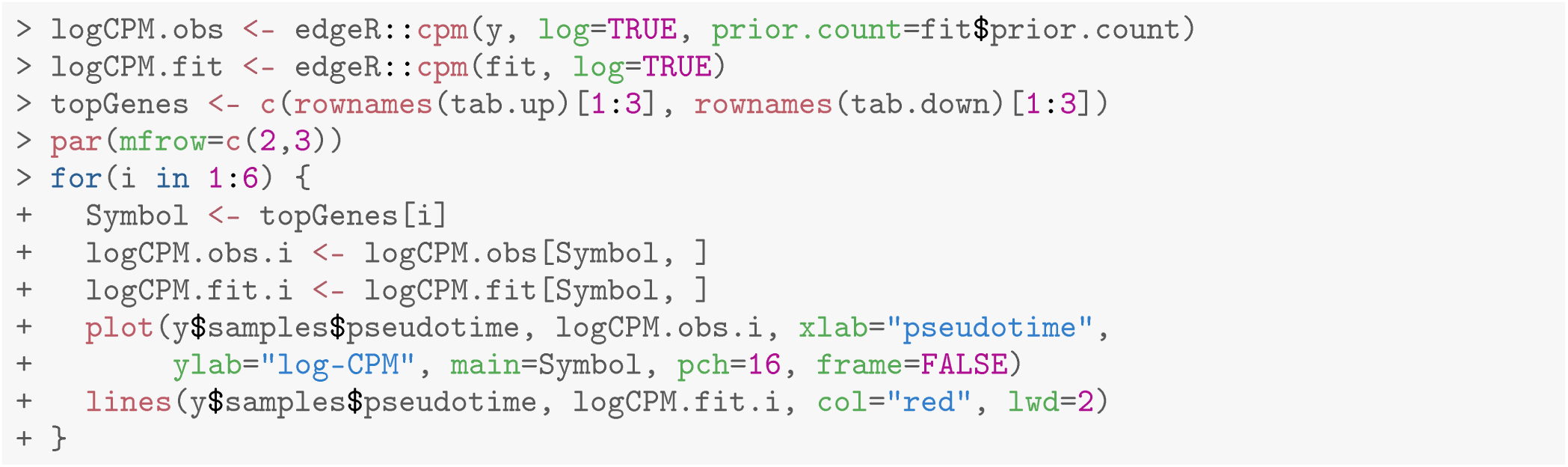

Scatter plots are produced to visualize the relationship between gene expression level and pseudotime for the top 3 increasing and the top 3 decreasing genes (Figure 10). Each point in the scatter plot indicates the observed `logCPM` of a pseudo-bulk sample at its average pseudotime. A smooth curve is drawn along pseudotime for each gene using the fitted `logCPM` values obtained from the spline model. The `cpm` function in *edgeR* is used to calculate the observed and fitted `logCPM`. Since there is a `cpm` function in the *SingleCellExperiment* package, we use `edgeR::cpm` to explicitly call the `cpm` function in *edgeR*. The smooth curves for the top row’s 3 genes exhibit a generally increasing trend in gene expression over pseudotime, while the curves for the bottom row’s 3 genes show a general decreasing trend.

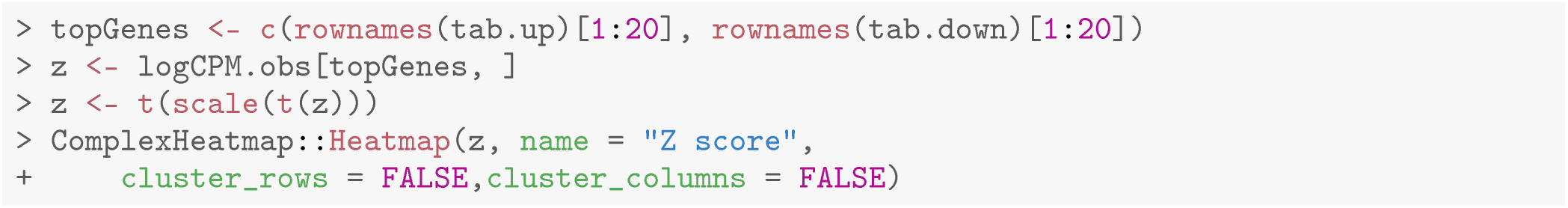

**Figure 10.**
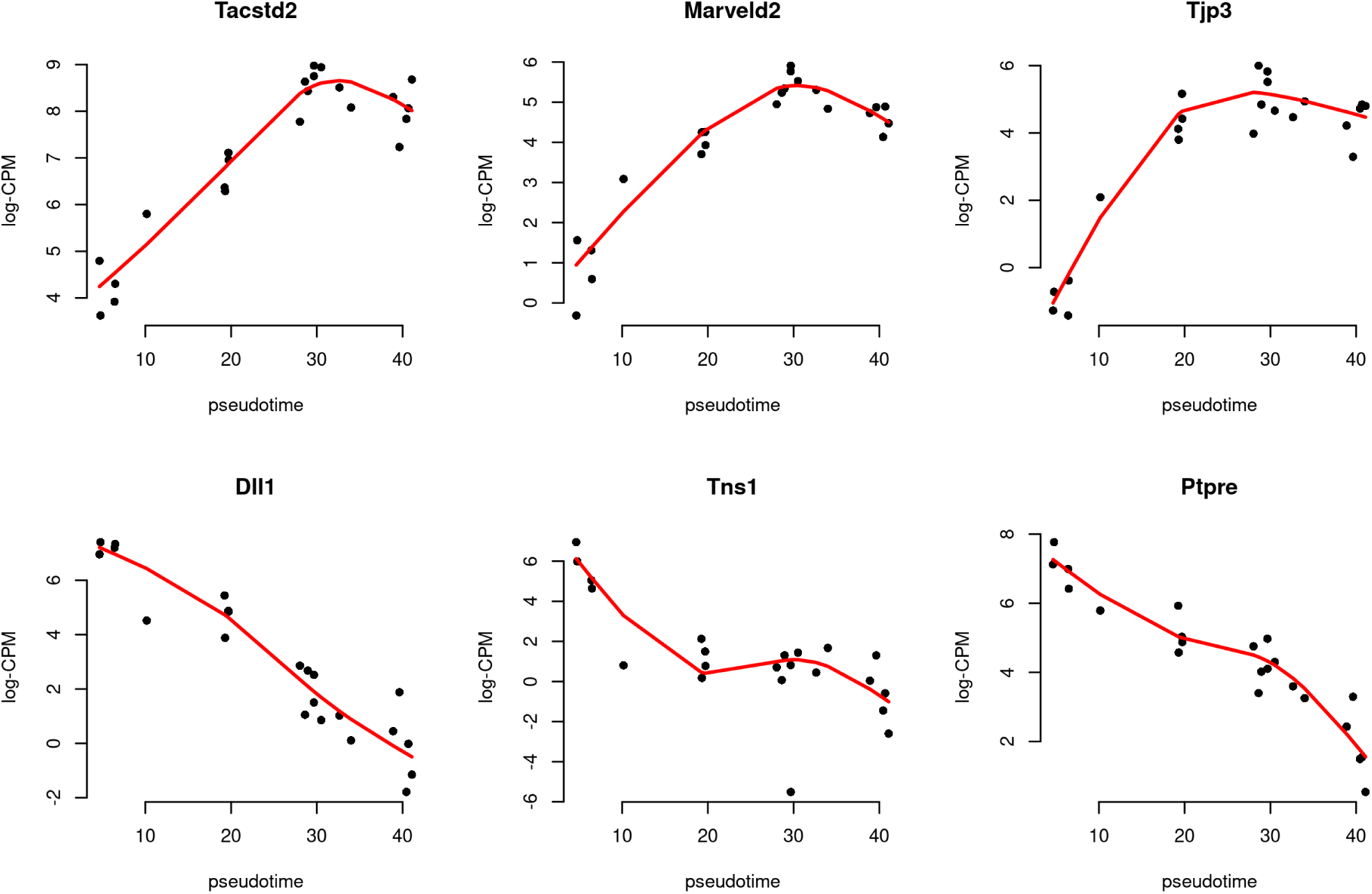
Scatter plots of expression of top genes along pseudotime. The black dots indicate the observed values, while the red line represents the fitted values calculated along pseudotime.

A heatmap is generated to examine the top 20 up and top 20 down genes collectively (Figure 11). In the heatmap, pseudo-bulk samples are arranged in increasing pseudotime from left to right. The up genes are on the top half of the heatmap whereas the down genes are on the bottom half. The heatmap shows a gradual increase in expression levels of the up genes from left to right, while the down genes display the opposite trend.

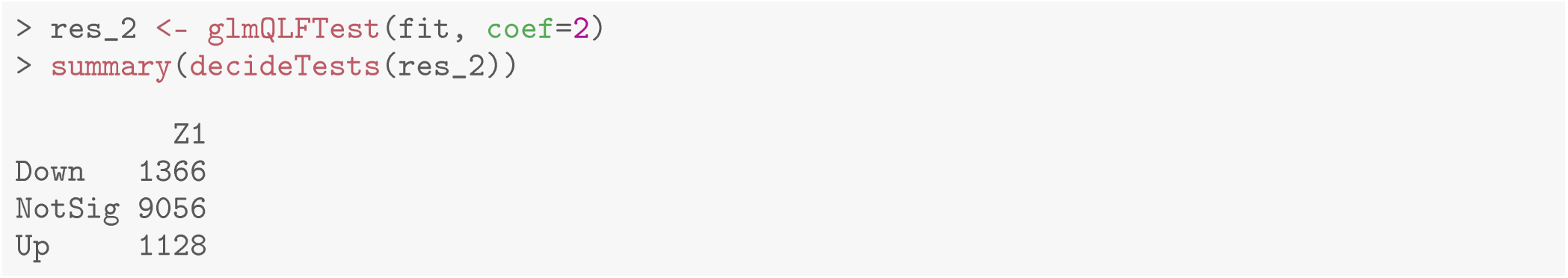

**Figure 11.**
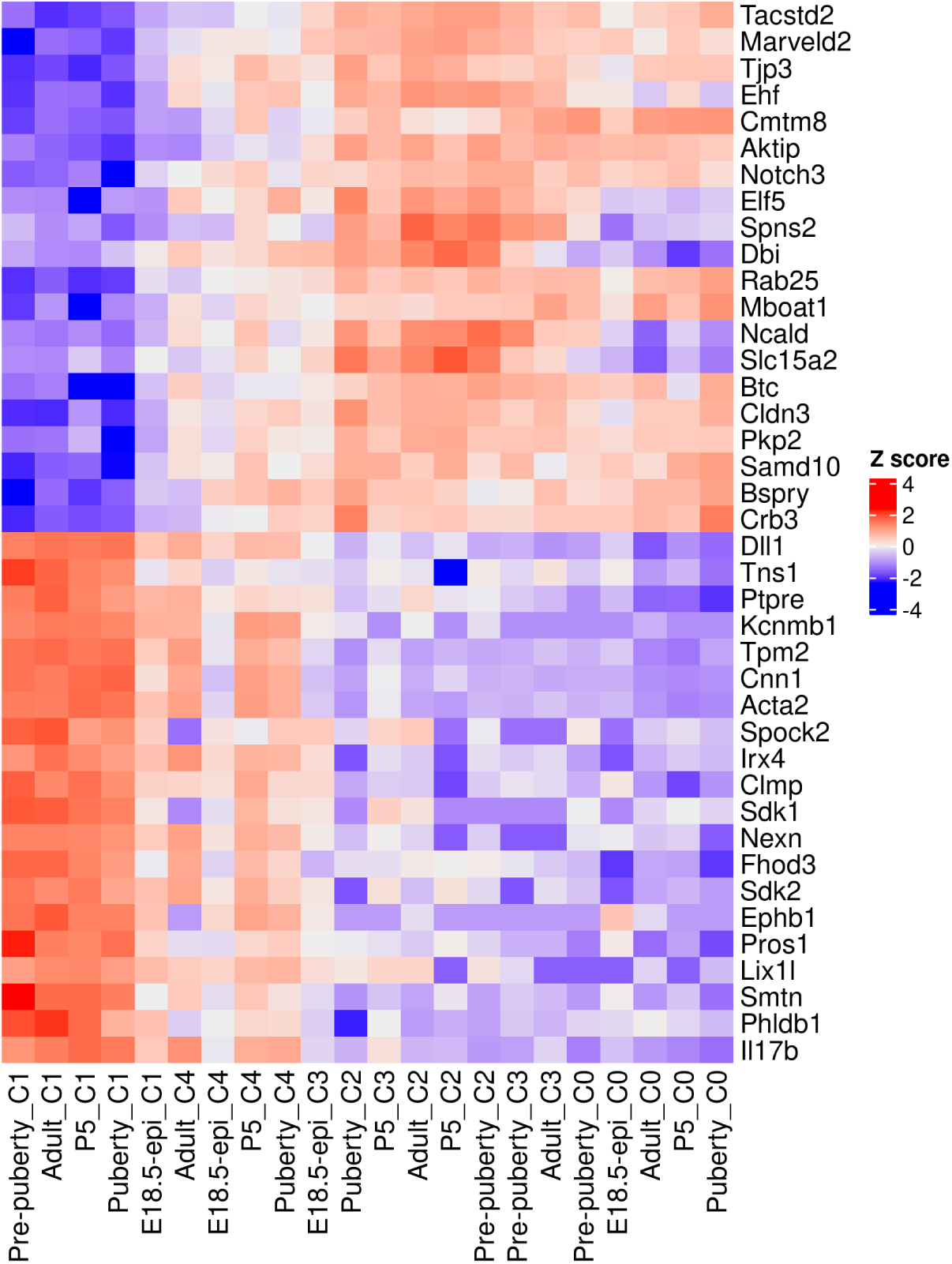
Heatmap of top 20 up and top 20 down genes. Rows are genes and columns are pseudo-bulk samples.

## Time course functional enrichment analysis

### Gene ontology analysis

To interpret the results of the time course analysis at the functional level, we perform geneset enrichment analysis. Gene ontology (GO) is one of the commonly used databases for this purpose. The GO terms in the GO databases are categorized into three classes: biological process (BP), cellular component (CC) and molecular function (MF). In a GO analysis, we are interested in finding GO terms that are over-represented or enriched with significant genes.

GO analysis is usually directional. For simplicity, we re-perform the QL F-test on the `Z1` coefficient to identify genes that exhibit a general linear increase or decrease along pseudotime. The numbers of genes with a significant increasing or decreasing linear trend are shown below.

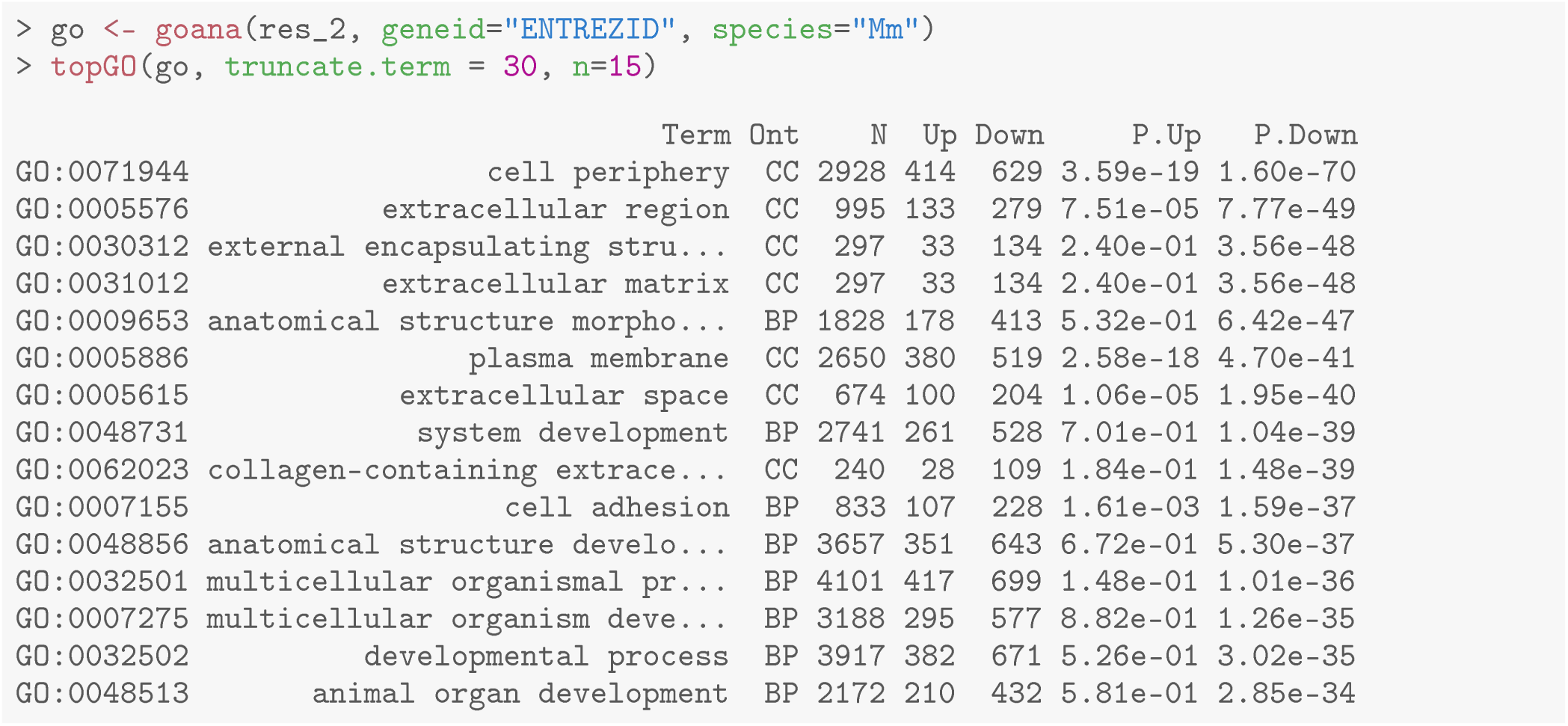

To perform a GO analysis, we apply the `goana` function to the above test results. Note that Entrez gene IDs are required for `goana`, which has been added to the `ENTREZiD` column in the gene annotation. The top enriched GO terms can be viewed using `topGO` function.

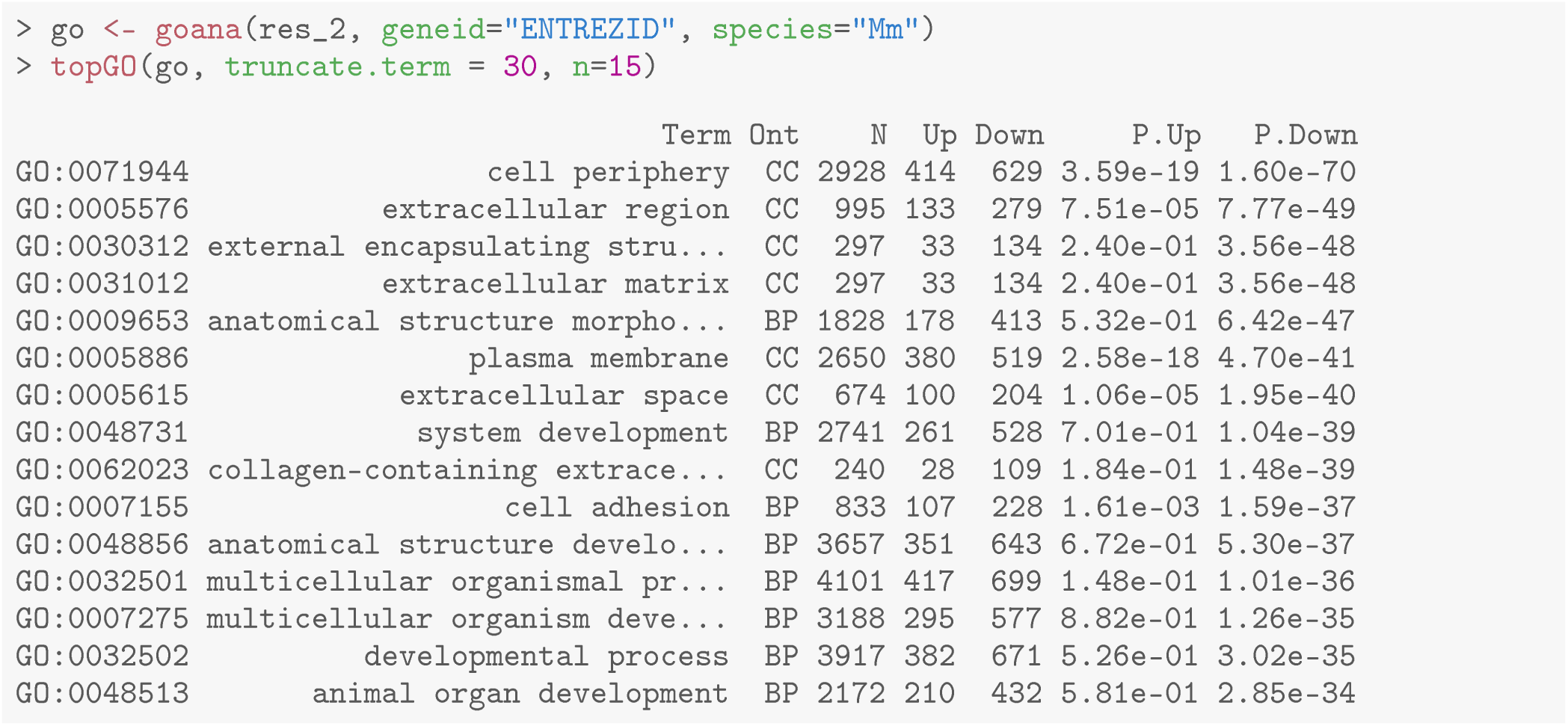

It can be seen that most of the top GO terms are down-regulated. Here, we choose the top 10 down-regulated terms for each GO category and show the results in a barplot (Figure 12).

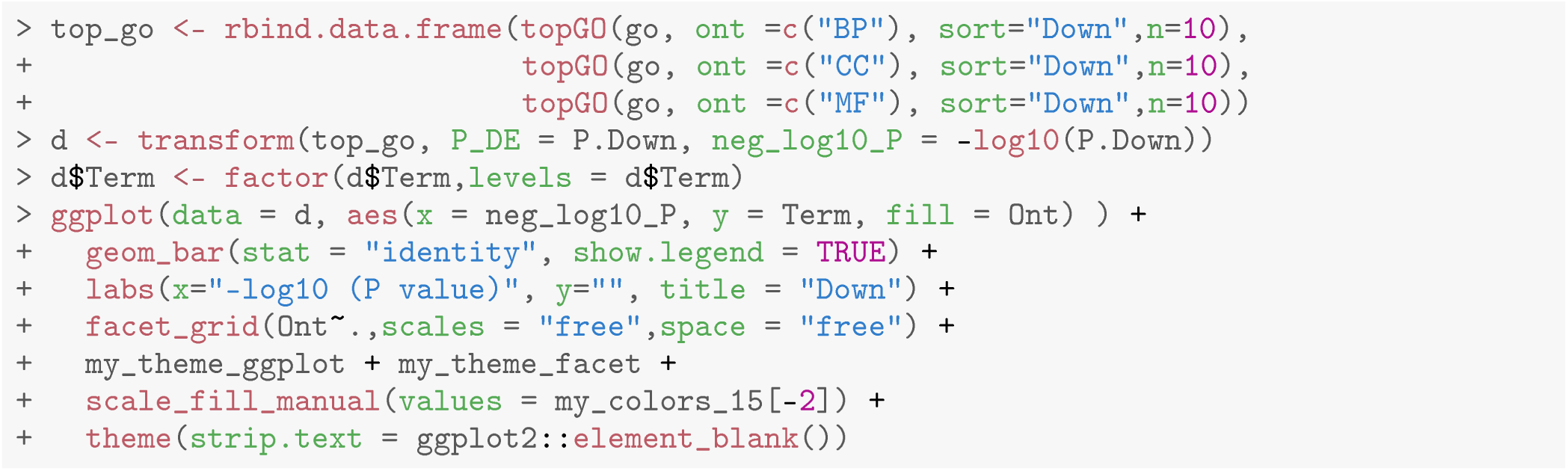

**Figure 12.**
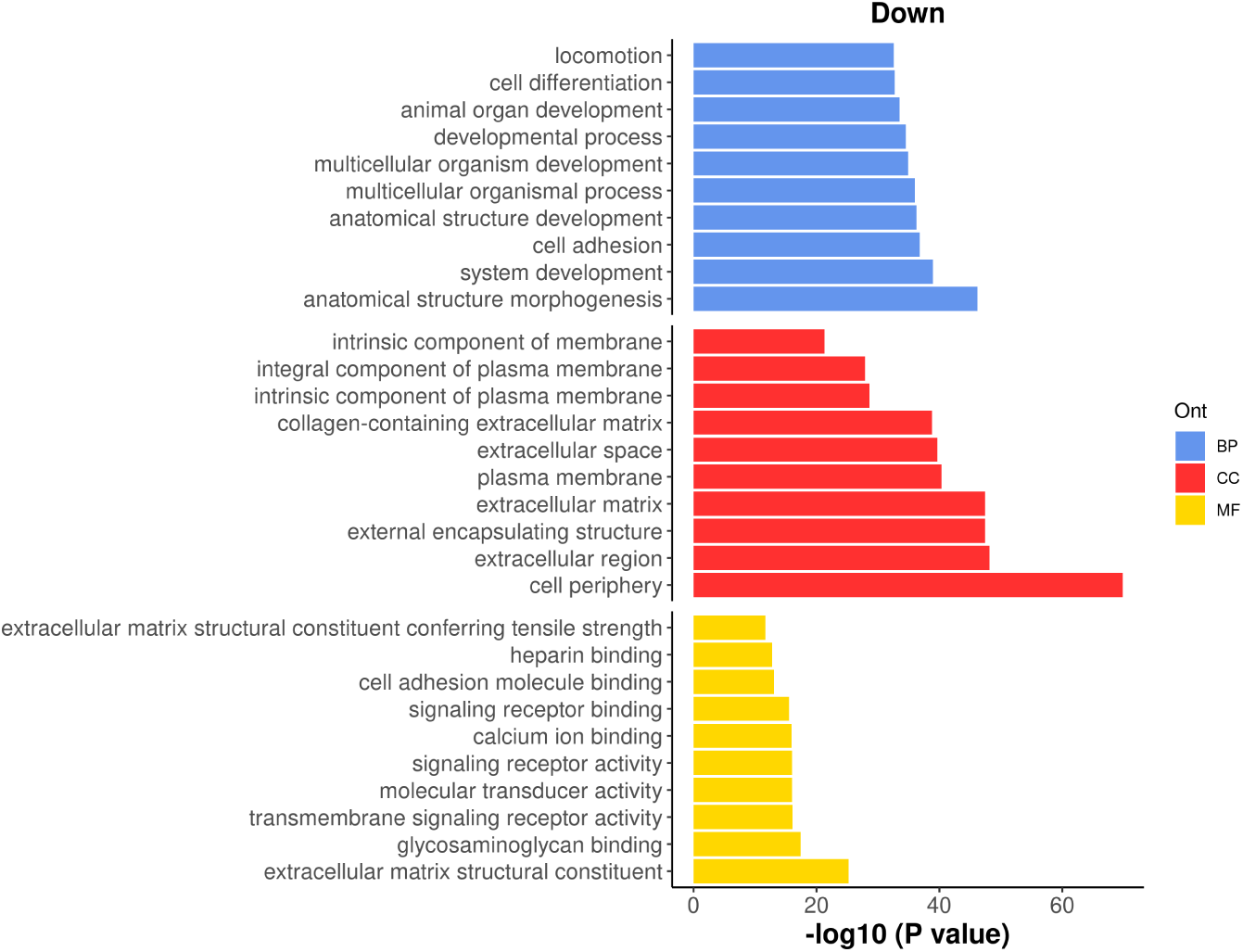
Barplot of *−* log_10_ p-values of the top 10 down-regulated GO terms under each GO category.

### KEGG pathway analysis

The Kyoto Encyclopedia of Genes and Genomes (KEGG)[23] is another commonly used database for exploring signaling pathways to understand the molecular mechanism of diseases and biological processes. A KEGG analysis can be done by using `kegga` function.

The top enriched KEGG pathways can be viewed by using `topKEGG` function.

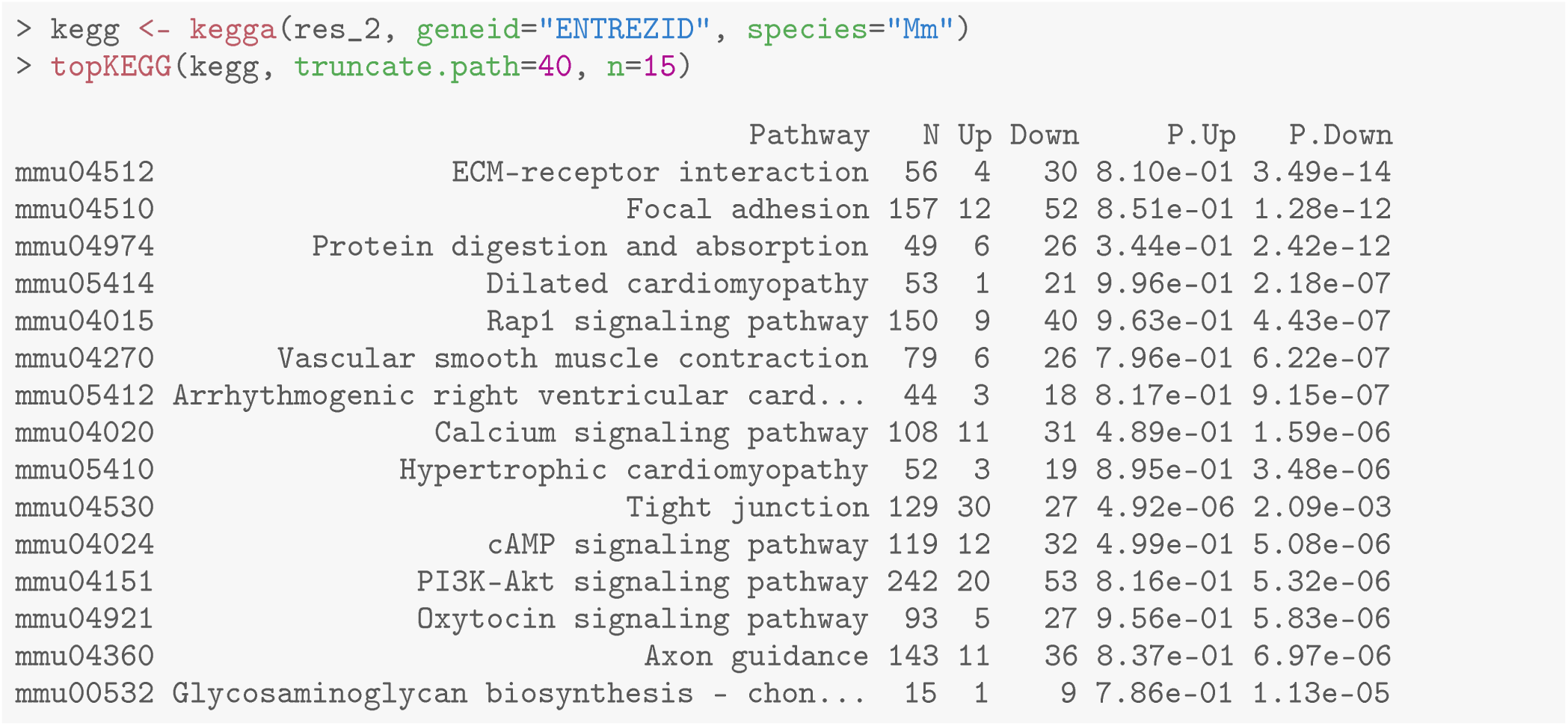

The results show that most of the top enriched KEGG pathways are down-regulated. Here, we select the top 15 downregulated KEGG pathways and visualize their significance in a barplot (Figure 13).

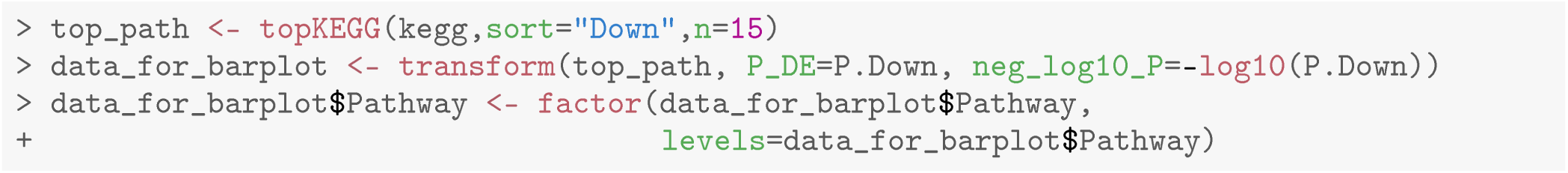

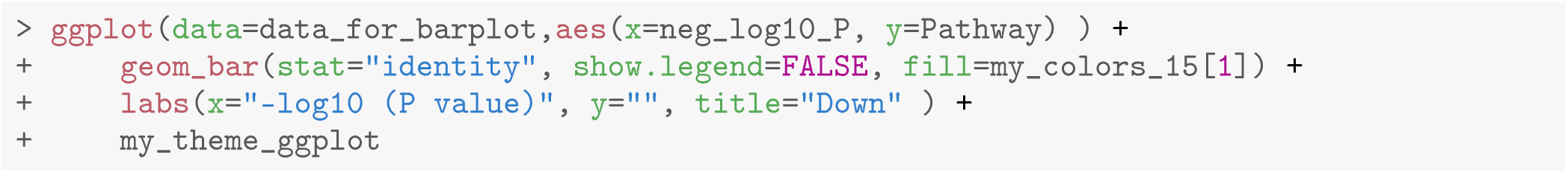

**Figure 13.**
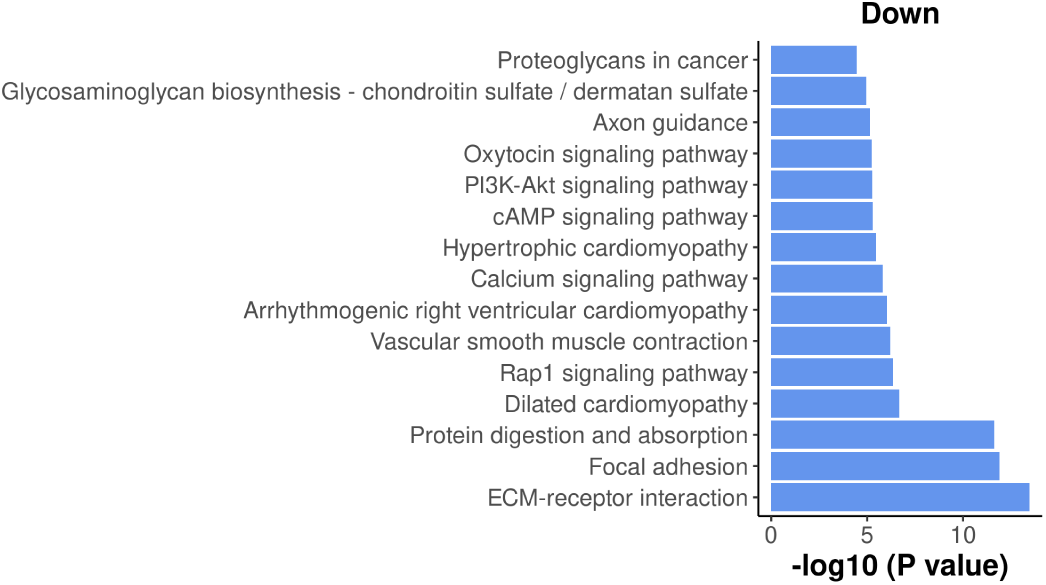
Barplot of *−* log_10_ p-values of the top 15 down-regulated KEGG pathways.

Among the top down-regulated pathways, the PI3K-Akt signaling pathway is noteworthy as it is typically involved in cell proliferation and plays a crucial role in mammary gland development.

To assess the overall expression level of the PI3K-Akt signaling pathway across pseudotime, a plot is generated by plotting the average expression level of all the genes in the pathway against pseudotime. The information of all the genes in the pathway can be obtained by `getGeneKEGGLinks` and `getKEGGPathwayNames`.

The plot below clearly illustrates a significant down-regulation of the PI3K-Akt pathway along pseudotime (Figure 14).

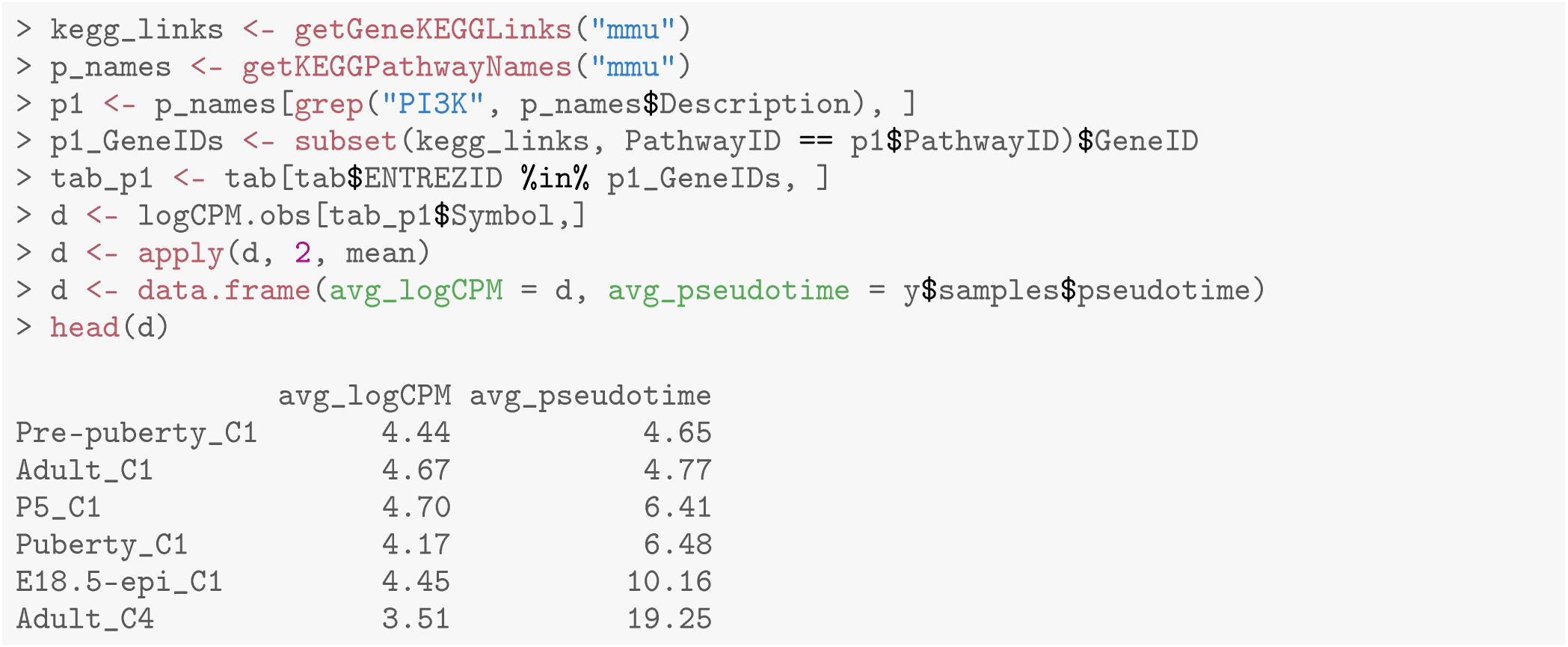

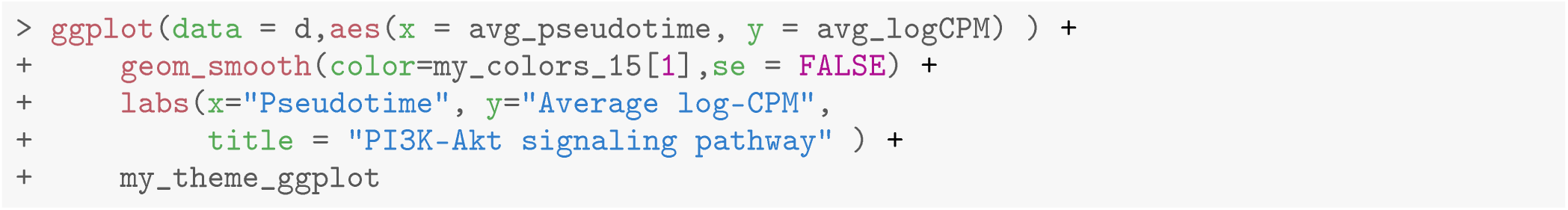

**Figure 14.**
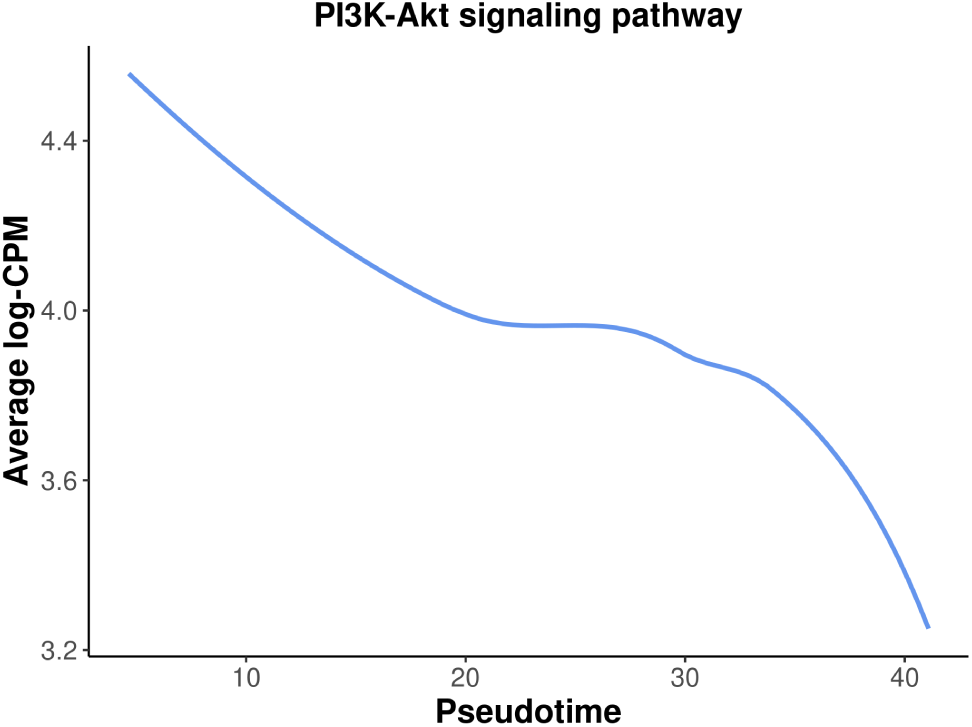
A smooth curve of PI3K-Akt signaling pathway expression level against spseudotime.

## Discussion

In this article, we demonstrated a complete workflow of a pseudo-temporal trajectory analysis of scRNA-seq data. This workflow takes single-cell count matrices as input and leverages the Seurat pipeline for standard scRNA-seq analysis, including quality control, normalization, and integration. The *scDblFinder* package is utilized for doublet prediction. Trajectory inference is conducted with *monocle3*, while the *edgeR* QL framework with a pseudo-bulking strategy is applied for pseudo-time course analysis. Alternative methods and packages can be used interchangeably with the ones implemented in this study, as long as they perform equivalent functions. For instance, the bioconductor workflow may be substituted for the Seurat pipeline in scRNA-seq analysis, whereas the *slingshot* package may replace *monocle3* for performing trajectory analysis.

This workflow article utilized 10x scRNA-seq data from five distinct stages of mouse mammary gland development, with a focus on the lineage progression of epithelial cells. By performing a time course analysis based on pseudotime along the developmental trajectory, we successfully identified genes and pathways that exhibit differential expression patterns over the course of pseudotime. The results of this extensive analysis not only confirm previous findings in the literature regarding the mouse mammary gland epithelium, but also reveal new insights specific to the early developmental stages of the mammary gland. The analytical framework presented here can be utilized for any single-cell experiments aimed at studying dynamic changes along a specific path, whether it involves cell differentiation or the development of cell types.

## Packages used

This workflow depends on various packages from the Bioconductor project version 3.15 and the Comprehensive R Archive Network (CRAN), running on R version 4.2.1 or higher. The complete list of the packages used for this workflow are shown below:

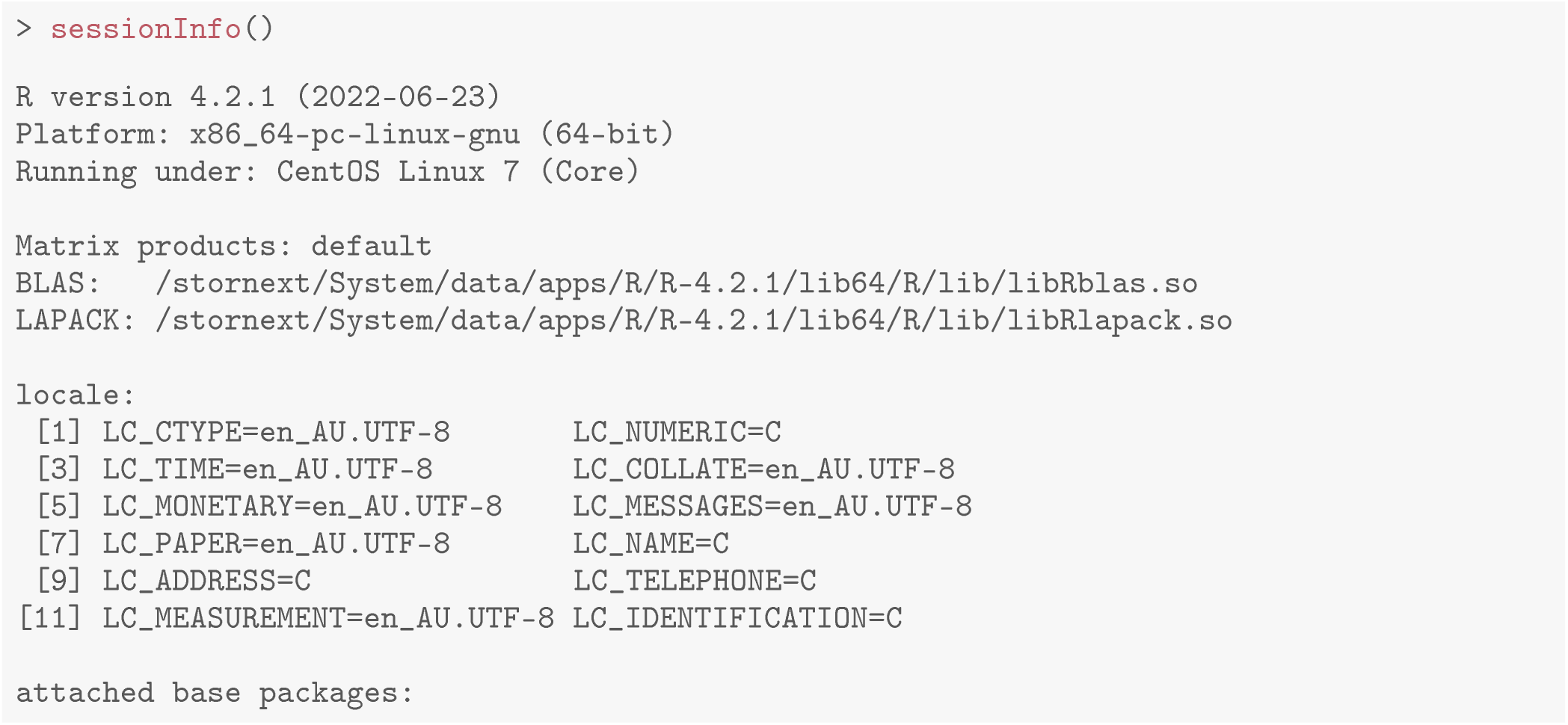

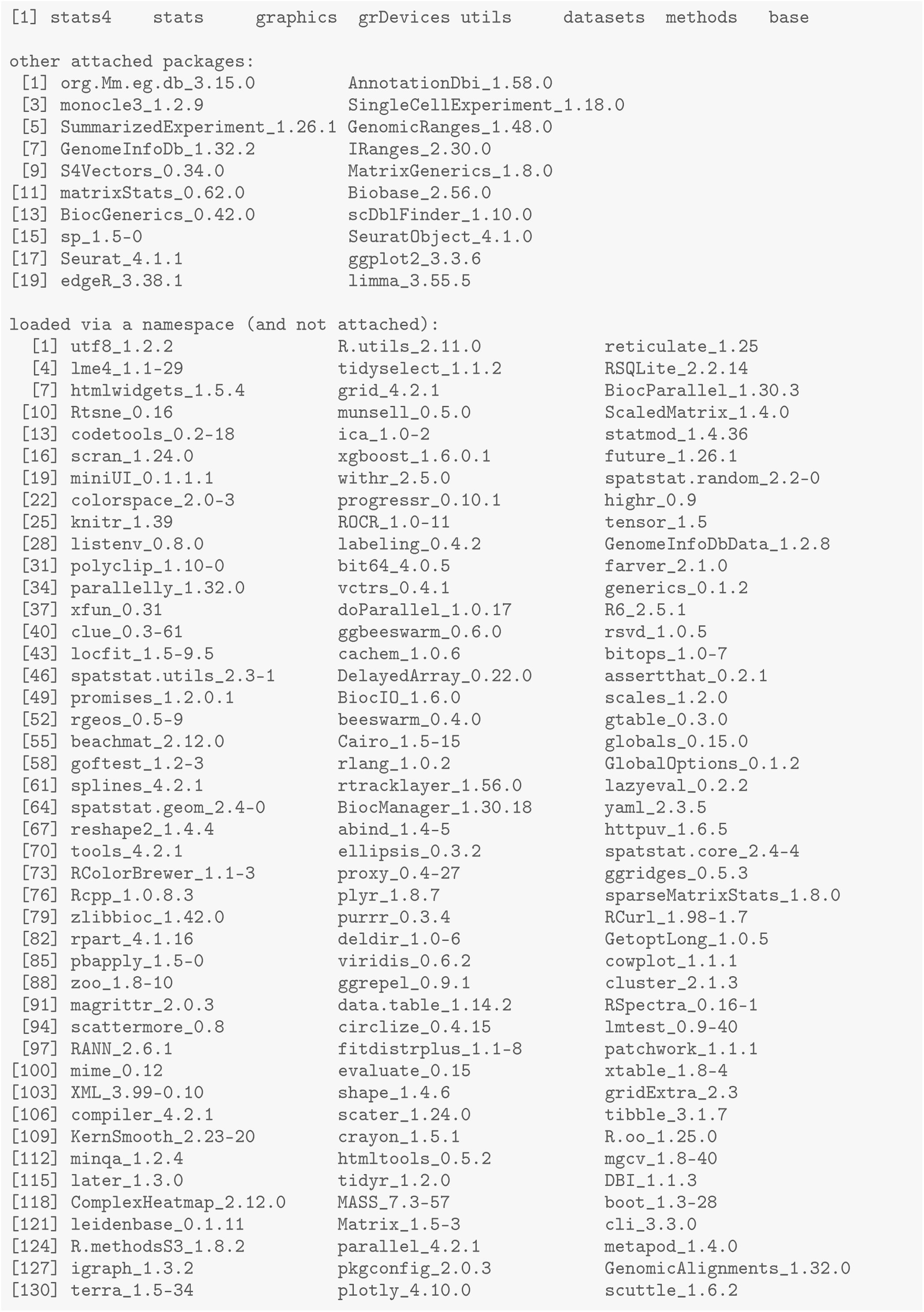

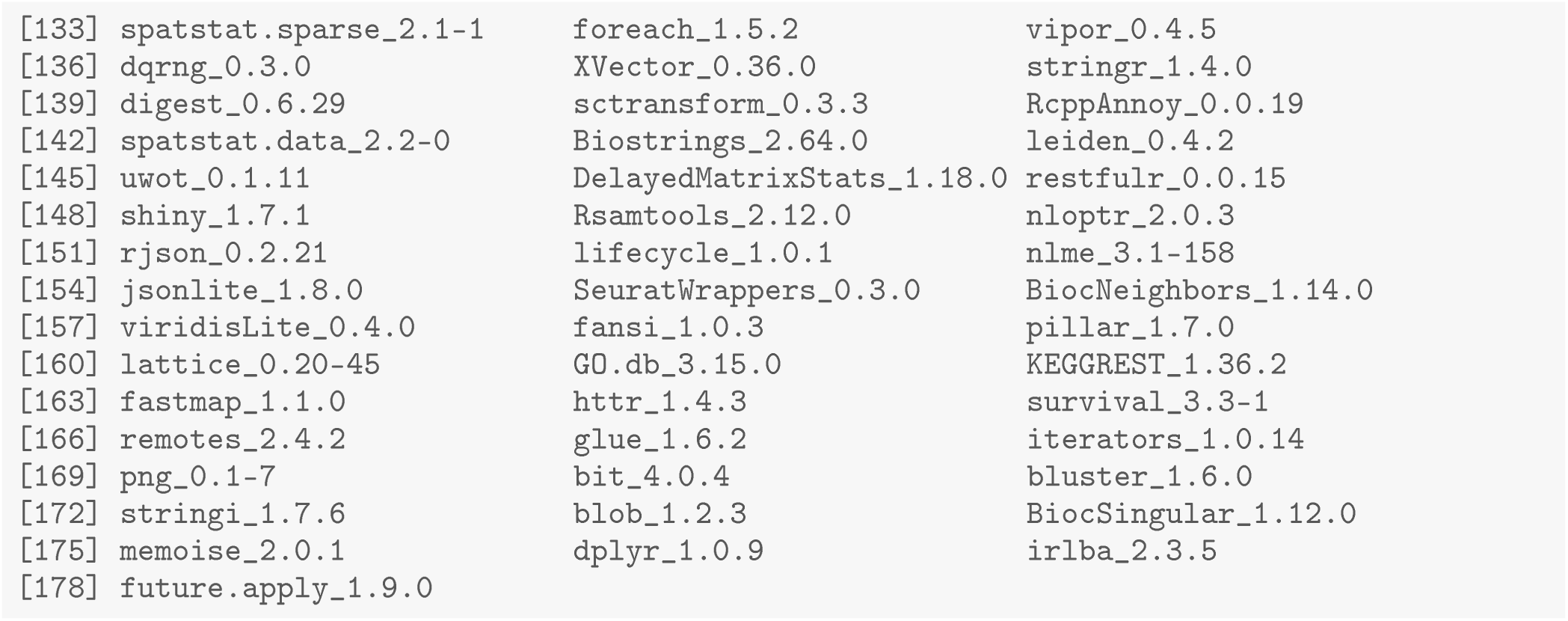

## Data availability

The single cell RNA-seq datasets used in this study were obtained from the Gene Expression Omnibus (GEO) with accession numbers of GSE103275 [16] and GSE164017 [17].

## Software availability

All the packages used in this workflow are publicly available from the Bioconductor project (version 3.15) and the Comprehensive R Archive Network (CRAN). The *knitr* Rnw file and R code that generated this article is available from: https://github.com/jinming-cheng/TimeCoursePaperWorkflow.

